# Crystal and cryo-EM structures of the cytosolic G protein alpha chaperone and guanine nucleotide exchange factor Ric-8A bound to Gαi1

**DOI:** 10.1101/2020.01.04.893156

**Authors:** Levi J. McClelland, Kaiming Zhang, Tung-Chung Mou, Jake Johnston, Cindee Yates-Hansen, Shanshan Li, Celestine J. Thomas, Tzanko I. Doukov, Sarah Triest, Alexandre Wohlkonig, Gregory G. Tall, Jan Steyaert, Wah Chiu, Stephen R. Sprang

## Abstract

Ric-8A is a cytosolic Guanine Nucleotide exchange Factor (GEF) that activates heterotrimeric G protein alpha subunits (Gα)^1^. Ric-8A is essential to life in multicellular eukaryotes by virtue of its chaperone activity that is required for Gα biogenesis and membrane localization^2, 3^. Ric-8A adopts an armadillo (ARM)/HEAT repeat domain architecture and is structurally unrelated to G Protein-Coupled Receptors (GPCR)^4^. Both GEF and chaperone activities are stimulated by Casein Kinase II phosphorylation^5^. The mechanisms by which Ric-8A catalyzes GDP release and GTP binding to Gα, or exerts chaperone activity are unknown. Here, we report the structure of the nanobody-stabilized complex of nucleotide-free Gαi1 (isoform 1 of Gα family i) and phosphorylated Ric-8A at near atomic resolution by cryo-electron microscopy and X-ray crystallography. We find that Ric-8A envelops the GTPase domain of Gα, disrupting all three switch regions that convey Gα nucleotide-binding and signaling activity, and displaces the C-terminal helix and helical domain of Gα. These cooperative interactions dismantle the GDP binding site and promote GDP release, while protecting structural elements of Gα that are dynamic in the nucleotide-free state. The structures also show how *in vivo* phosphorylation stabilizes Gα-binding elements of Ric-8A, thereby enhancing its GEF and chaperone activities.

---

Genetic^6–8^ and biochemical data^9^ support a role for Ric-8A in G protein-coupled receptor (GPCR)-independent regulation of asymmetric cell division that is essential for embryonic development^10^. Ric-8A exhibits GEF and chaperone activity towards G protein alpha subunits (Gα) of the I, q and 12/13 classes^1^, while Ric-8B, performs these functions for Gαs and Gαolf - each in a variety of cellular contexts^11, 12^. That the Gα-class specificity of Ric-8A and Ric-8B is the same for GEF and chaperone activities^13^ suggests a common mechanistic basis for both.

Biophysical data show that nucleotide-free Gαi1 is structurally dynamic when bound to Ric-8A^14, 15^ and shares some properties with GPCR-bound Gα^16^. Namely, Ric-8A induces rotational dynamics in the Gα helical domain^17^ and binds to the C-terminus of Gα ^14, 18^. In contrast to GPCRs, there is evidence that Ric-8A forms extensive interactions with the Gα switch regions that undergo GTP-dependent conformational changes^1, 15, 18, 19^ and with the Gα β-sheet scaffold^15^.

To better understand the structural basis of Ric-8A GEF and chaperone activity, we have determined the structure of the Ric-8A:Gαi1 complex, using both cryo-electron microscopy (cryo-EM) and X-ray crystallography. To form the complex we used the N-terminal 491-residue fragment of rat Ric-8A, which is a more active GEF then the full-length (530 amino acid) protein, and, to reduce conformational heterogeneity, we used the N-terminal 31-residue truncation mutant of rat Gαi1 (Δ31NGαi1), an efficacious substrate for Ric-8A^14^. Recombinant Ric-8A(1-491) was phosphorylated by Casein Kinase II at S435 and T440, which is necessary and sufficient for stimulation of GEF activity^5^ **(Extended data Fig. 1)**. Hereafter we refer to phosphorylated Ric-8A(1-491) as Ric-8A and Δ31NGαi1 as Gα.

**Figure 1.**
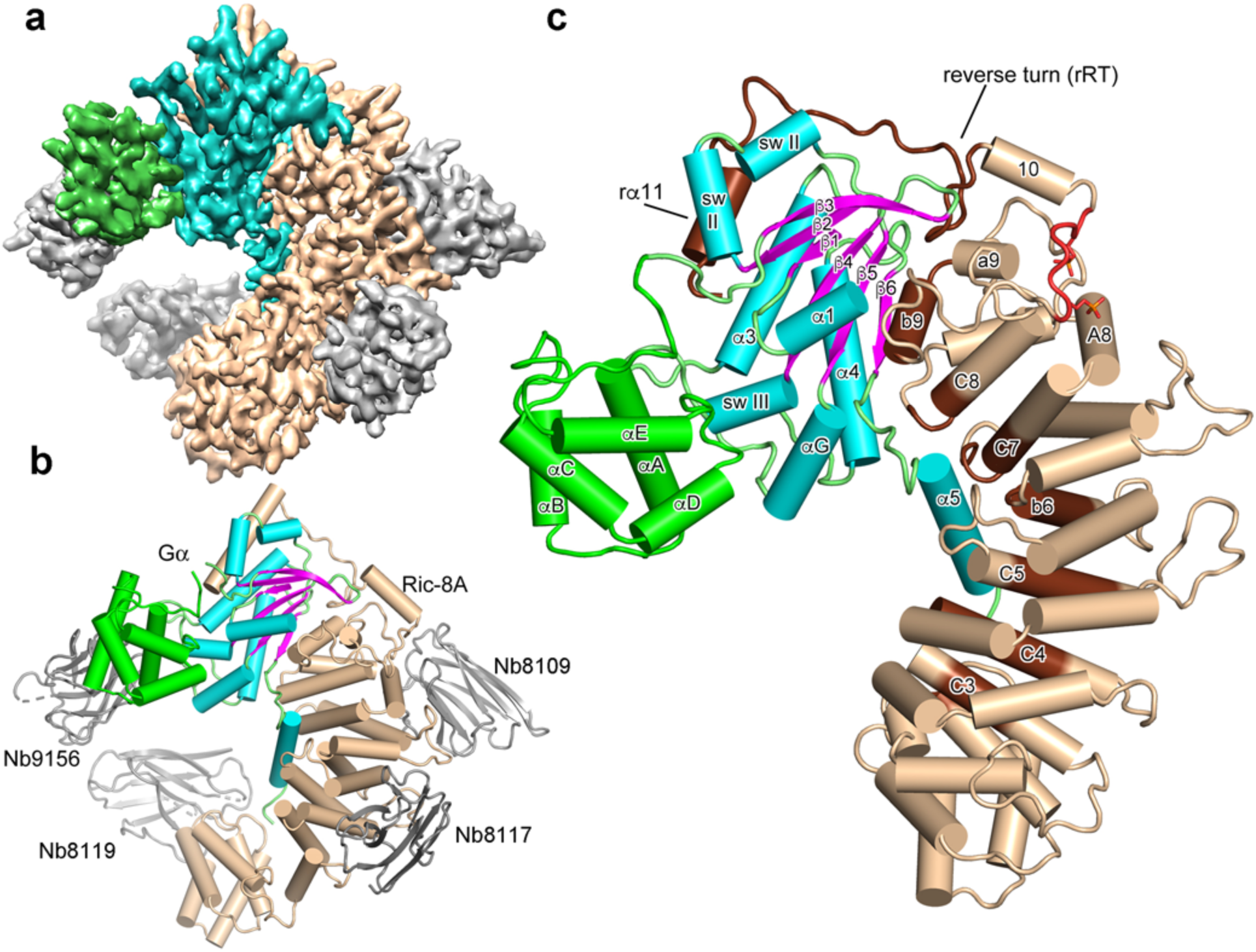
Architecture of the Ric-8A:Gα:4Nb complex. **a**, Cryo-EM 3D reconstruction of Ric-8A:Gα:4Nb is shown with Ric-8A colored *wheat*, Gα GTPase and Helical domains colored *cyan* and *green*, respectively, and nanobodies colored *gray*. **b**, Annotated ribbon and cylinder drawing, colored as in **a**, but with the β-strands of Gα rendered in *magenta*. (**C**) The Ric-8A:Gαi1:4Nb complex is shown with the nanobodies removed and the loop segments of the Gα GTPase domain colored green. Segments of Ric-8A that contact Gα are rendered in *dark brown*. Ric-8A residues 335-340, which include the two phosphorylation sites are rendered in *red*.

To stabilize, and limit the dynamics of the Ric-8A:Gα complex for crystallographic and cryo-EM experiments, we developed a panel of camelid nanobodies (Nb)^20^ that specifically recognize either Gαi1, Ric-8A or the complex of the two. We formed a series of Ric-8A:Gαi1:Nb complexes that included from one to four Nbs from this panel. The stability of the complexes towards cryogenic vitrification and consequently, the quality and resolution of cryo-EM reconstructions derived from them, was improved in step with the number of Nbs in the complex. We determined the cryo-EM structure of a complex of Ric-8A:Gαi1 with three Nbs bound to Ric-8A and one bound to the helical domain of Gα.

Together, these nanobodies do not significantly affect Ric-8A-GEF activity (**Extended Data Fig. 2**). The structure was determined from 327,493 particles derived from 8,468 movie images, yielding a 3.9Å resolution reconstruction (**Fig. 1, Extended Data Figs. 3** and **4 and Table 1**).

**Figure 2.**
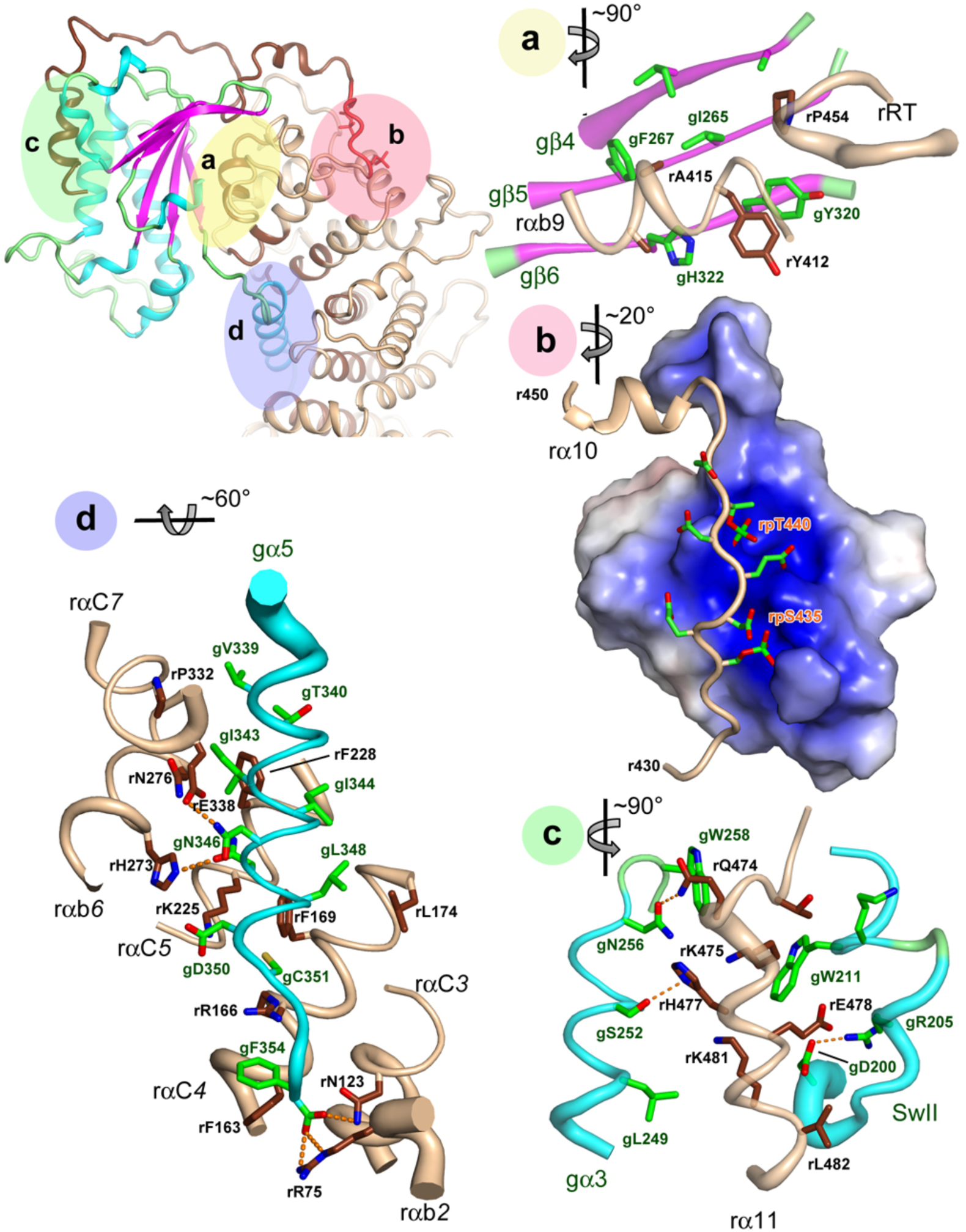
Interactions of Ric-8A with Gαi1. Three major Ric-8A:Gα contact surfaces and phosphorylation sites are highlighted and labeled in the *green*, *yello*w, *pink* and *blue* overlays (upper left) and enlarged in panels **a**-**d**. Axes adjacent to each panel label indicate the rotation applied to the overall schematic to generate the panel view. **a,** Interaction between Gα β-sheet and the Ric-8A C-terminal ARM/HEAT repeat helix rαA*9* and reverse turn r451-r457 (rRT) from the crystal structure. Polypeptide backbones are rendered as tubes, with diameter proportional to B-factor at Cα. Carbon atoms of Ric-8A and Gαi1 are rendered in *deep brown*, and green, respectively, and oxygen and nitrogen atoms are respectively colored *red* and *blue*. **b,** Acidic Ric-8A peptide with phosphorylated residues rpS335 and rpT440 bound to the positively charged surface formed by the 8^th^ and 9th Ric-8A ARM/HEAT repeats from the crystal structure. **c,** Interaction of rα11 with gα3 and Switch II from cryo-EM. Putative hydrogen bonds (donor-acceptor contact < 3.5Å) are depicted with *orange* dashed lines. **d,** Contacts between gα5 and residues in successive ARM/HEAT repeats of Ric-8A, from the crystal structure.

**Figure 3.**
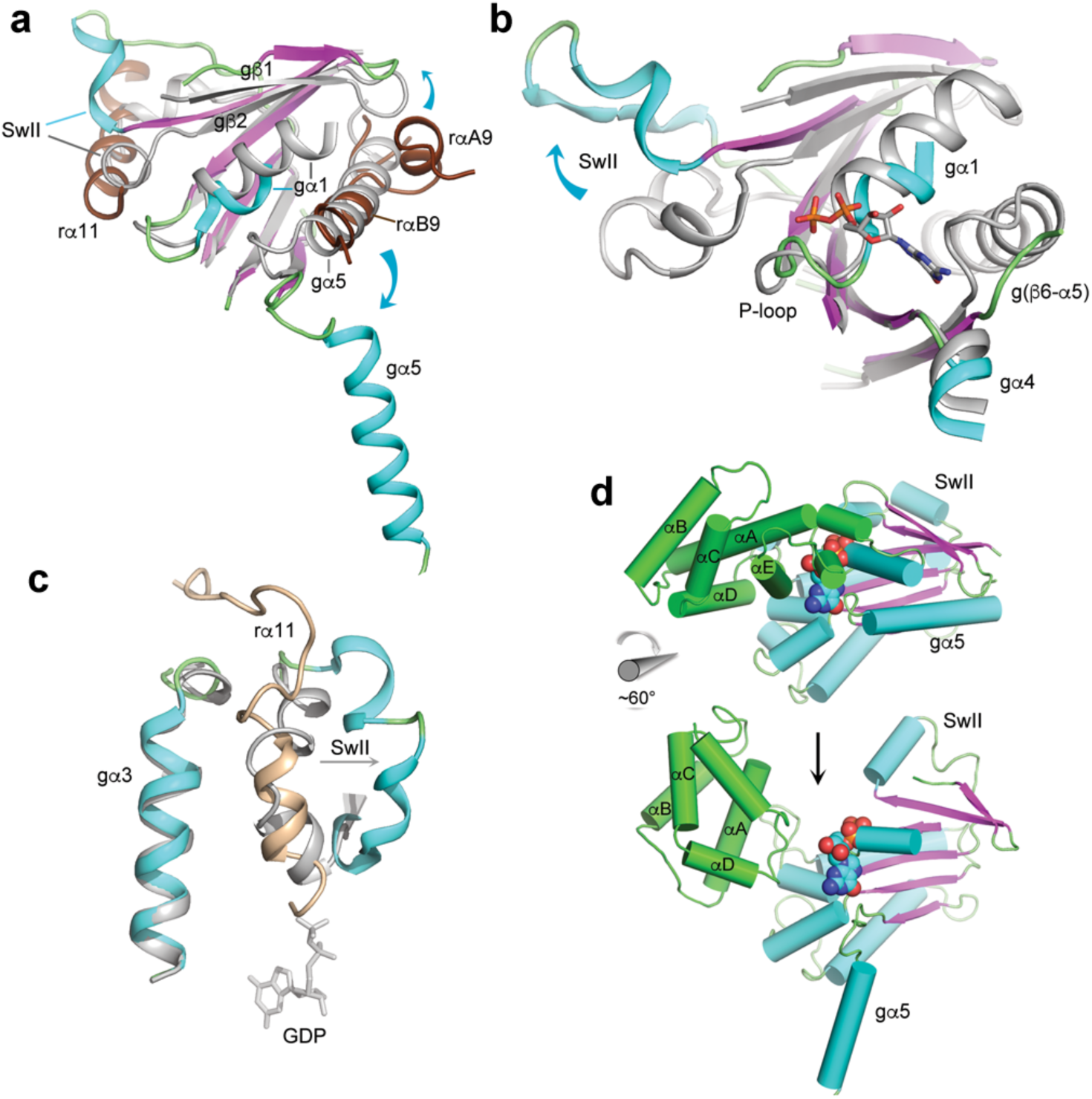
Conformational changes induced in Gα by its interaction with Ric-8A. In panels **a-c**, GDP•Pi-bound Gαi1 (PDB ID 1GIT), rendered in *gray* is superimposed on the crystal structure of Ric-8A-bound Gα using the Cα atoms of Gαi1 residues 219-225 (gβ4), 262-270 (gβ5) and 318-325. **a,** Conformational changes due to binding of Ric-8A αa9, αb9 and RT to Gαi1. **b,** Ric-8A-induced conformation changes dismantle the Gα nucleotide binding site (see text). GDP from 1GIT is included as a stick model for reference. **c**, Displacement of Switch II by rα*11*. The position of GDP bound to Gαi1•GDP is shown as a stick model. **d,** Ric-8A-induced rotation of the helical domain away from the GTPase domain of Gα: *top*, Gαi1•GDP (1GIT) rendered with helices as cylinders and β-strands as ribbons and the helical domain colored green; atoms of GDP are rendered as spheres. *Bottom*, the cryo-EM – derived model of Ric-8A-bound Gα with GDP from 1GIT shown as a reference point.

**Figure 4.**
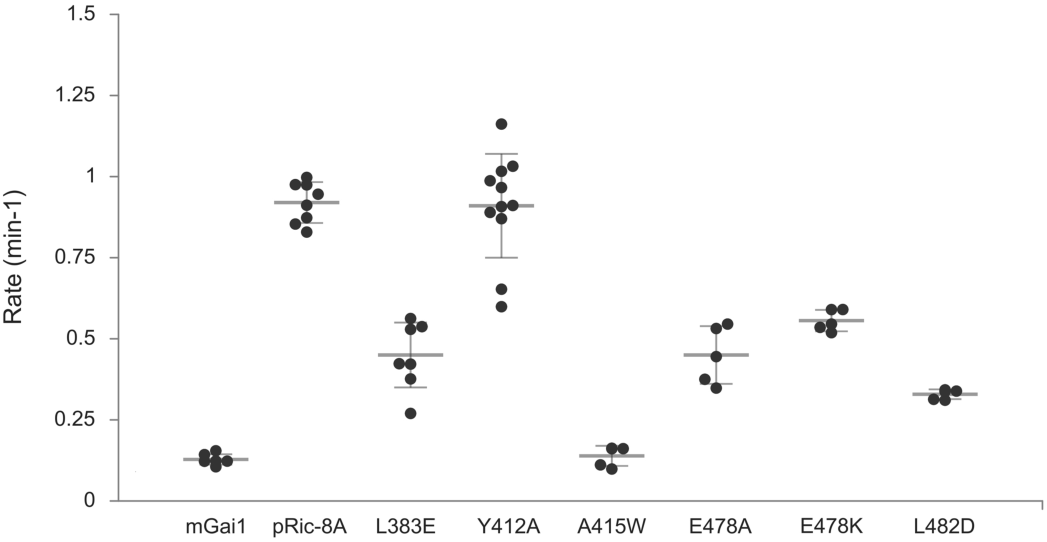
Mutational analysis of selected residues at the Ric-8A:Gα interface. GDP-GTP exchange rates were measured by the rate of tryptophan fluorescence increase upon addition of ΔN31Gαi1 to Ric-8A and GTPγS at final concentrations of 2μM Ric-8A, 1μM ΔN31Gαi1 and 10μM GTPγS. Rate differences between wild-type Ric-8A and all of the Ric-8A mutants with the exception of Y412A (P<0.82), are significant at 10^-5^ > p >10^-12^. Horizontal bars represent means and 1 standard deviation.

We obtained crystals of Ric-8A:Gαi1 bound to the three Ric-8A-specific Nbs used to generate the cryo-EM structure. Diffraction from these crystals was highly anisotropic (**Extended Data Fig. 5**), extending to 4.6Å along *a\**, and *b*,* and 3.3Å along *c\**, affording measurement of a 90% complete anisotropic dataset to 3.3Å resolution **(Extended Data Table 2**). Initial crystallographic phases were determined by molecular replacement and subsequently used to fit the cryo-EM density map. Iterative cycles of model-building and refinement utilized both cryo-EM and X-ray diffraction data to generate the final models **(Extended Data Table S1 and S2)**. In the following discussion, we use prefixes “r” and “g” for residue and secondary structure identifiers of Ric-8A and Gα, respectively. ARM/HEAT repeat helices of Ric-8A are designated according to their position in the repeat: “A”, “B” or “C” for ARM repeats or “a” and “b” for HEAT repeats, followed by the sequence number of the repeat (1 through 9). We use the established nomenclature for Gα secondary structure ^21–23^. Descriptions of the crystal structure of Ric-8A:Gα refer to chains A and B, the better ordered of the two complexes in the asymmetric unit.

**Figure 5.**
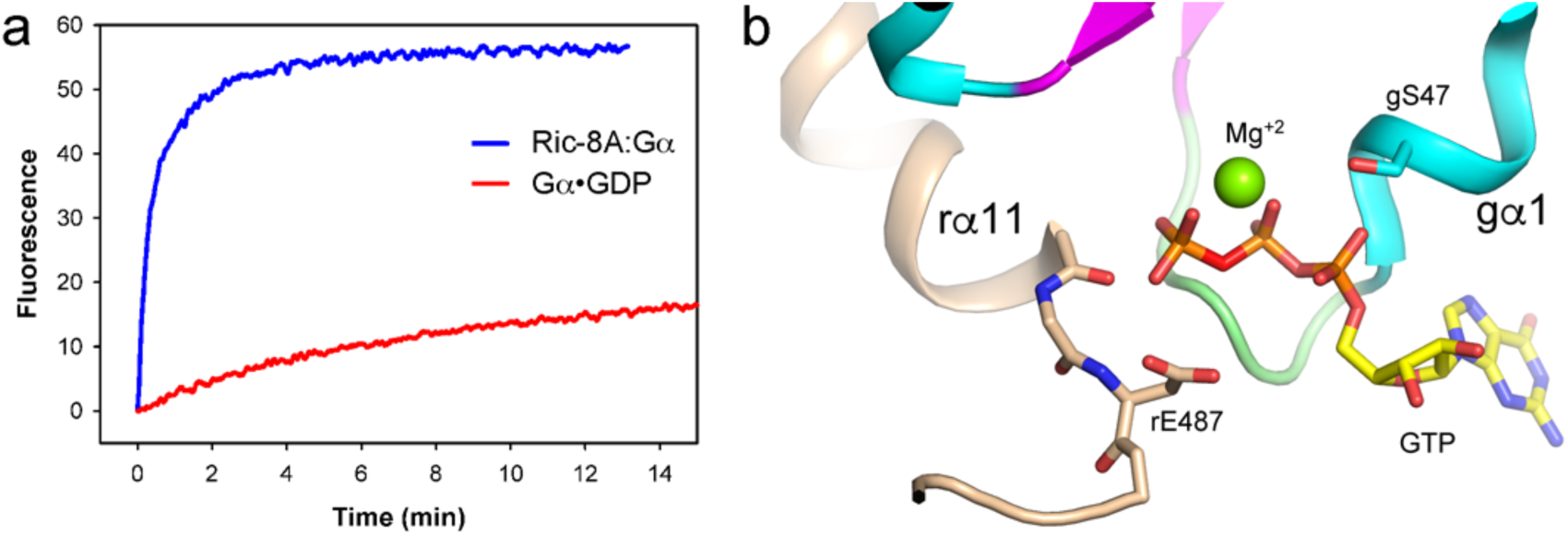
The P-loop of Gαi1 provides a pre-ordered binding site for GTP in the complex with Ric-8A. **a**, Progress curve for GTP (10μM) binding to purified Ric-8A:Gαi1 complex (1μM) (*red*), relative to that for nucleotide exchange at Gαi1•GDP (1μM). **b**, Superposition of a model of GTP (PDB ID 1CIP) onto the nucleotide binding site of Ric-8A-bound Gαi1. Serine 47 of α1 is positioned for Mg^+2^ coordination (see text).

The cryo-EM reconstruction reveals Ric-8A residues 2-487 and the whole of Gα with the exception of the disordered linker (residues 50-76) between the Helical and GTPase domains (**Fig. 1a** and **1b and Extended Data Video 1**). The crystal structure of the Ric-8A:Gα complex also reveals continuous electron density for Ric-8A except for the linkage between 422 and 430. Notably, residues C-terminal to the last HEAT repeat (r430-r491) are disordered in structures of Ric-8A in which Gα is absent^4, 18^. In the crystal structure, which lacks Nb1056, the helical domain of Gαi1 and its connections to the GTPase domain, including much of gα1 and all of switch I, are disordered.

Otherwise the cryo-EM and X-ray models of Ric-8A:Gα are in good agreement, although certain irregularly-structured and loop regions in both Gα and Ric-8A show significant divergence (**Extended Data Fig. 6**). Small angle X-ray scattering (SAXS) measurements of the complex are consistent with the X-ray and cryo-EM structures (**Extended Data Fig. 7**).

Binding of Ric-8A to the GTPase domain of Gα at three non-contiguous contact surfaces bury more than 3200Å^2^ of solvent-accessible surface area (**Fig. 2a,c,d**). Together, these interactions destabilize the guanine nucleotide binding site (**Fig. 3**). At the core of the complex, Ric-8A αb9 (r401-r408) and the reverse turn r451-r457 (rRT) interact with the gβ4-gβ6 strands of Gα (**Fig. 2a and Extended Data Figs 8a** and **b**). Ric-8 αb9 occupies the site of gα5 in GDP-bound Gα, and consequently gα5 is ejected from the concave surface of the Gα β-sheet (**Fig. 3a**). Ric-8A residues rY412, rA415, rA416 and rL418 in rαb9 substitute for nonpolar residues of gα5 to stabilize the hydrophobic surface of the Gα β-sheet core. These residues are conserved in both A and B isoforms of vertebrate Ric-8 (**Extended Data Fig. 9a**) and form Van der Waals interactions with residues in gβ4-gβ6 that are conserved in both Gαi and Gαs families (**Extended Data Fig. 9b**). The rW415A mutation severely impairs GEF activity (**Fig. 4**), although the Y412A mutation does not. At the same interface, steric interactions with rαa9 lever the antiparallel gβ2-gβ3 hairpin away from the GTPase core (**Fig. 3a**), thereby destabilizing and partially disordering gα1. This, as described below, triggers separation of the Helical and GTPase domains of Gα.

The ∼90° rotation of α5 away from the GTPase domain core (**Fig. 3a**) reconfigures the “TATC” motif in the g(β6-α5) GDP purine binding loop (**Fig. 3b)**. This structural change perturbs the conserved NKKD motif (in gβ4-αG) that confers specificity for guanosine nucleotides^24^. Displacement of these two loops dismantles the GDP purine binding site. The hydrophobic contact between the N-terminus of gα4 and the Ric-8A αB8-αC8 turn (**Extended Data Fig. 10)** is also important, as indicated by the impairment of GEF activity resulting from the rL383E mutation (**Fig. 4**).

After its ejection from the Gα β−sheet, gα5 is accommodated in a broad trough formed by helices rαb2 through rαB8 that line the concave surface of ARM/Heat superhelix of Ric-8A (**Fig 2d and Extended Data Fig. 8g** and **h**), as observed also in the structure of Ric-8A bound to the C-terminus of transducin^18^. The predominantly hydrophobic amino acid residues of gα5 that interact with the Ric-8A trough (e.g. gF336, gV339, gI343, gI344 and gL348) otherwise pack against the GTPase domain β-sheet of nucleotide-bound Gα^25^. Most of the Ric-8A residues that contact gα5 are conserved in Ric-8B **(Extended Data Fig. 9a**), whereas several of the Ric-8A-contacting residues in gα5 are not conserved among Gα classes (**Extended Data Fig. 9b**).

Ric-8A phosphorylation acts as an entropic clamp to promote GEF activity. The Ric-8A αb9 and RT segments that interact with the Gα β-sheet are connected by an intrinsically disordered sequence, r430-r450^4^ followed by rα10. The segment from r430-r440 is rich in acidic amino acids, and binds within a basic groove formed by rαA8 (r344-r358) and rαa9 (r401-r410) (**Fig. 2b**). Phosphorylated rpS435 and rpT440, which are well-ordered (**Extended Data Fig. 8i-l**) and form multiple ion-pair interactions with conserved lysine and arginine side chains in the Ric-8A electropositive groove, help to immobilize the r430-450 connector and consequently stabilize rαb9 and rRT that interact with gβ4-gβ6. Accordingly, we found earlier that chargeàneutral mutations of rR345Q and rK349A, which interact with rpS435 and adjacent acidic residues, reduce both basal and phosphorylation-stimulated GEF activity of Ric-8A^4^.

Ric-8A helix α11 (r471-r491), which is preceded by an elongated peptide (r458-r470) that forms an extended arch over the α3-β5 loop of Gα, packs between Gα Switch II and gα3 (**Fig. 2c, and Extended Data Fig 8c-f**). In the GTP-bound state of Gα, these two elements form the Gα effector protein binding site^26^. rα11 occupies the position of Switch II in the G protein heterotrimer^27^, and also in Gαi1 bound to GTP analogs^25^ and in the Gαi1•GDP•Pi product complex^28^ (**Fig. 3c**). However, Switch II is disordered in Gαi1•GDP and probably also in nucleotide-free Gα^22^. Ric-8A interactions with switch II are functionally important, since mutations of two Gα-contacting residues in rα11, rE478 and L482, which are conserved in both Ric-8A isoforms, impair GEF activity (**Fig. 4**). The conformations of Switch II observed in the X-ray and cryo-EM structures differ from each other, and from those adopted in nucleotide-bound Gα^26^ (**Extended Data Fig. 8e,f**). Remodeling of Switch II may be facilitated by Ric-8A-induced displacement of β2-β3 (**Fig. 3a**) described above.

Displacement of gα5 and reorientation of gβ2-gβ3 eliminates a nexus of stabilizing interactions with the C-terminal residues of gα1, which become disordered (**Fig 3a**.). As a consequence, contacts between the GTPase and helical domains of Gα are disrupted. As observed in the cryo-EM structure, the helical domain undergoes a clockwise rotation of ∼60° around an axis roughly aligned with αD of the Gα helical domain (**Fig. 3d**). By this motion, a channel opens between the helical and GTPase domains, providing a path of egress for the nucleotide. The magnitude of the rotation is less than that observed in crystal and cryo-EM structures of GPCR:G protein complexes^16, 29^, or deduced from double electron-electron resonance studies of Ric-8A:Gα^17^, possibly because it is limited by steric interactions between the nanobodies bound to Gα and Ric-8A. The variability in the orientation of the helical domain is also suggested by normal-mode analysis of the SAXS data (**Extended Data Fig. 7**). The coupled interactions between Ric-8A and Gαi1, described above, induce profound conformational changes that dismantle the Gα guanine nucleotide-binding site (**Fig. 3b and Extended Data Video 2**).

The basis for the Gα class selectivity of Ric-8 isoforms is not readily apparent. The great majority of residues at the Gα contact residues are conserved in Ric-8A and Ric-8B (**Extended Data Fig 9a**). Gα residues that interact with rαB9 and rRT are conserved in the Gαi and Gαs classes. However, several residues in gα5 differ between Gαs and Gαi classes (**Extended Data Fig. 9b**), and, as suggested^18^, the latter may engage in more productive interactions with Ric-8A than their Gαs counterparts.

The structure of the complex suggests elements of Ric-8A chaperone activity by which it promotes folding and stabilization of nucleotide-free Gα^2^. It is remarkable that the magnitude of conformational changes induced in Gαi1 by Ric-8A far exceed those sufficient to effect GDP release in GPCR - G protein complexes^16, 23^ wherein gα5 undergoes a modest rotation and translation and switch II remains protected by Gβγ as in the intact heterotrimer^27, 30^. Rather, Ric-8A stabilizes nucleotide-free Gα subunits in the absence of Gβγ^14^. Molecular dynamics simulations suggest that contacts between gα5 and the Gα β-sheet are dynamic in nucleotide-free Gα^31^. Interactions with Ric-8A would shield these hydrophobic surfaces from exposure in the cytosol. Switch II is also a dynamic structure, even in the GDP-bound state of Gα^24^ and would likewise be stabilized by Ric-8A α11. Ric-8A(1-491) partially rescues Gαi1 biosynthesis in Ric-8A ^-/-^ cells, but not that of Gαq, for which full-length Ric-8A is required^32^. Hence, the C-terminal ∼40 residues of Ric-8A, which harbor three of the five CKII phosphorylation sites^5^, must play a critical role in Ric-8A chaperone activity. Nevertheless, it is clear that the mechanism by which Ric-8A stabilizes dynamic regions of the nucleotide-free Gα GTPase domain also underlie its GEF activity.

The extensive interface between Ric-8A and Gα prompts the question how the two proteins are able to dissociate upon binding of GTP to Gα, a process that is kinetically facile in comparison to the overall exchange reaction (**Fig. 5a and Extended Data Fig. 2**). Although displaced slightly by steric conflict with rα11 (**Fig. 3b**), the conformation of the P-loop is largely retained in the nucleotide-free Gα bound to Ric-8A (**Fig. 5b**), thus providing a preformed platform for subsequent binding of GTP. Electrostatic repulsion between the C-terminus of r11 and the γ-phosphate of GTP as modeled in **Fig. 5b**, could promote its release from its binding site between gα3 and Switch II, allowing Switch II to refold into its native GTP-bound conformation. Disruption of Switch II interactions with Ric-8A would restore the native structure of gβ2-gβ3, promote its interaction with gα1 and thereby destabilize the remaining interface of Ric-8A with the GTPase domain. The dynamics that accompany Ric-8A binding to Gα•GDP and subsequent release of Gα•GTP remain to be explored.

## Methods

### Preparation of crosslinked Ric-8A and Gαi1

Bis-sulphosuccinimidyl suberate (BS3), Pierce-Thermo-Fisher Scientific) was dissolved in water to a concentration of 100 mM. K100 reagent (CovalX) is provided as a 2 mg/ml aqueous solution and contained a proprietary mixture of inert carbon chain spacers (lengths between 8.8-13.2 Å), separating 1-hydroxyl-7-azabenzotriozole groups. Ric-8A:Gαi1 complex, prepared as described^1–3^ was dialyzed into 20 mM phosphate buffer saline (PBS) pH 7.4, 1 mM DTT, and final concentration adjusted to 20 mM in the same buffer. For each 100 ml of complex, 20 ml of K100 (2mg/ml) and 25 ml of BS3 (100 mM) was added, incubated at room temperature for 30 minutes and the reaction quenched by addition of 10 ml of 1 M Tris pH 8.5. Samples were centrifuged at 14,000 rpm on a desk-top centrifuge for 10 min at 4°C, to remove particulate matter, eluted though a tandem 5 ml HiTrap desalting column (GE Healthcare) to remove excess cross-linking reagents. Proteins were eluted in 20 mM PBS, concentrated to 0.3 mg/ml and 1 mM fresh DTT added as the final step. Cross-linked samples were analyzed by Coomassie-stained SDS-PAGE and verified by Western blotting using an anti-Ric-8A monoclonal antibody^4^ and mouse Anti-Gαi1 monocolonal antibody (Enzo Lifesciences). The crosslinked preparation contained cross-linked Ric-8A:Gαi1, free Gαi1 and Ric-8A in roughly equal proportion.

### Development of Camelid nanobodies

Llamas (*Lama glama*) were immunized with the cross-linked Ric-8A:Gαi1 preparation. Peripheral blood lymphocytes were isolated from the immunized animals for extraction of total RNA, generation of cDNA and PCR amplification to isolate cDNAs encoding heavy chain variable domains (VHH) for subcloning into the pCTCON2 vector for display on the cell surface of *Saccharomyces cerevisiae* EBY100 cells, as described^5–7^. Cells expressing nanobodies that selectively recognize Gαi1, Ric-8A or the crosslinked Ric-8A:Gαi1 were identified by three-color FACS sorting on a FACS AriaIII (BD Biosciences), using Dylight 505 (Pierce, Thermo Scientific)-conjugated Ric-8A, Dylight 488-conjugated Gαi1 and R-phycoerythrin goat anti-mouse antibody (Lucron) to label nanobody-expressing clones. Three rounds of selection were conducted using 10mM fluorescent proteins in 20 mM PBS pH 7.4, 100mM NaCl and 2mM DTT. Nanobody sequences from selected clones yielding nanobodies that exclusively bind to Gαi1, to Ric-8A either free or bound to Gαi1, or exclusively to the Ric-8A:Gαi1 complex were subcloned in a pMESy4 vector for periplasmic expression in *E. coli* WK6 cells as described^8^. Nanobody sequences encode a C-terminal hexahistidine affinity tag.

Of the set of 27 nanobodies discovered in screening that show strong expression, twenty were capable of binding to either free Ric-8A or to Ric-8A in complex with Gαi1 but not to Gαi1, three bound exclusively to the Ric-8A:Gαi1 complex and three bound to Gαi1 or its complex with Ric-8A but not to Ric-8A alone. From this set was identified a set of four nanobodies capable of binding simultaneously to Ric-8A:Gαi1, as determined by co-elution as a stable hexameric complex from a GE Healthcare HiLoad 16/600 Superdex 200pg column and validated by mass spectrometry on a MicroFlex MALDI-ToF MS(Brucker Microflex). Of this set, Nb8109, Nb8117 and Nb8119 bind to Ric-8A and the Ric-8A:Gαi1 complex and Nb9156 binds to Gαi1.

### Nanobody expression and purification

Nanobodies (Nb) 8109, 8117, 8119 and 9156 were encoded in pMESy4 vectors for periplasmic expression in WK6 *Escherichia coli* cells^7^. Briefly, overnight cultures were grown in 50 ml LB media containing 100 µg/ml Ampicillin, 100 mM Glucose, and 1 mM MgCl_2_ in shaker flasks at 190 rpm and 37 °C. After ∼16 hours cells were pelleted at 4000 rpm (2200xg) for 10 minutes using a benchtop Sorvall Legend RT. Re-suspended pellets were added to 1 L TB media containing 100 µg/ml Ampicillin, 5 mM glucose, and 1 mM MgCl_2_ then incubated at 190 rpm and 37 °C until achieving an OD_600_ of 0.7-1.0, at which point temperature was lowered to 28 °C and growths were induced with 300 µM isopropyl β-D-1-thiogalactopyranoside (IPTG). Approximately 16 hours post-induction, cells were pelleted at 8000 rpm (12000xg) for 15 min in a Sorvall RC 6+ Centrifuge and cell pellets were stored at – 80 °C. Pellets were re-suspended in 100 ml of TES buffer (0.2 M Tris pH 8, 0.5 mM EDTA, 0.5 M sucrose) at 4 °C, vortexed and allowed to stir for a minimum of 1 hour. Cells were then added dropwise to a 200 ml solution of TES/4 buffer (0.05 M Tris pH 8, 0.125 mM EDTA, 0.125 M sucrose), then stirred for 1 hour at 4 °C. Lysate was then spun down at 8000 rpm (12000xg) for 30 min and supernatant was loaded on a gravity column with Ni-NTA agarose resin (Qiagen). Nanobody-bound resin was washed with 20 column volumes of Wash 1 (0.05 M Tris pH 8, 1 M NaCl), 5 column volumes of Wash 2 (0.05 M MES pH 6, 1 M NaCl), 10 column volumes of Wash 3 (0.05 M Tris pH 8, 0.5 M NaCl), and then eluted with 3 column volumes of Elution Buffer (0.05 M Tris pH 8, 0.2 M NaCl, 0.5 M imidazole). Eluted nanobody was dialyzed overnight in 0.05 M Tris pH 8 and 0.2 M NaCl, then concentrated to 5-10 mg/mL before use or storage at −80 °C. Before use all nanobodies were gel purified using a Superdex 200 10/300 GL size-exclusion column (GE Healthcare) in gel filtration buffer: 50 mM HEPES pH 8, 150 mM NaCl, and 1 mM TCEP.

Complexes of Ric-8A, ΔN31Gαi1 and nanobodies Nb8109, Nb8117, Nb8119 in the presence or absence of Nb9156 were formed by incubating Ric-8A, ΔN31Gαi1 and nanobodies at a 1:2:2(per nanobody) molar ratio on ice for 16 hours and subsequently purified by gel filtration chromatography using a GE Healthcare HiLoad 16/600 Superdex 200 pg in Gel Filtration Buffer.

### Gα and Ric-8A protein expression and purification

N-terminally glutathione-S-transferase tagged Rat Gαi1 with a 31 residue N-terminal truncation (ΔN31Gαi1) was expressed from a pDest15 vector in Bl21(DE3) RIPL *Escherichia coli* and purified as previously described^1, 9^. N-terminally hexahistidine tagged Rat Ric-8A(residues 1-491, hereafter Ric-8A) was expressed and purified as previously described in a pET28a vector with BL21(DE3) RIPL *Escherichia coli* cells^1, 3, 10^. Following removal of the hexahistidine tag using Tobacco Etch Virus protease, Ric-8A protein was loaded onto a Source 15Q column and eluted with a 500 mM NaCl gradient at 180 mM NaCl and subsequently loaded on a GE Healthcare HiLoad 16/600 Superdex 200 column for size exclusion chromatography in gel filtration buffer containing 50 mM Tris pH 8, 150 mM NaCl and 1 mM Tris(2-carboxyethyl)phosphine (TCEP).

Ric-8A was phosphorylated using casein kinase II (New England Biolabs) as previously described^10, 11^. Briefly, purified Ric-8A was mixed 1:1 with a 2X Kinase Reaction Buffer containing 0.1 M HEPES pH 8, 0.1 M NaCl, 20 mM MgCl_2_, 2 mM EGTA, and 1 mM DTT. ATP was added to a final concentration of 5 mM. 160 Units of casein kinase II were added per mg of Ric-8A, and the reaction was allowed to proceed for ∼16 hours at room temperature. Phosphorylated protein was loaded onto a Source 15Q column pre-equilibrated with 50 mM HEPES pH 8, 25 mM NaCl, and 2 mM β-ME, and eluted from a 500 mM NaCl gradient at 210 mM NaCl. Phosphorylated Ric-8A exhibits an approximately 2 mS/cm shift compared to un-phosphorylated Ric-8A (**Extended Data Fig. 1b**). All procedures described hereinafter were conducted with phosphorylated Ric-8A comprising residues 1-491 of the full-length protein and referred to as Ric-8A. Myristoylated Gαi1 (mGαi1), used in nucleotide exchange assays, was prepared as described previously^10^. Mutants of Ric-8A, used in nucleotide exchange assays, were constructed via the QuikChange II XL Site-Directed Mutagenesis Kit (Agilent). Expression, purification and phosphorylation proceeded as described above.

### Guanine Nucleotide Exchange assays

were conducted by measuring the change in Gαi1 tryptophan fluorescence in the presence or absence of Ric-8A using either ΔN31Gαi1 or mGαi1, as described^3^. All proteins were buffer exchanged and assays were conducted in 50 mM HEPES pH 8, 150 mM NaCl, 10 mM MgCl_2_, 1 mM TCEP. Experiments were carried out at 20°C using a LS55 luminescence spectrometer (Perkin Elmer) with 4 nm slit widths (excitation 295 nm, emission 345 nm). GEF activity for Ric-8A and Ric-8A mutants were measured by preincubating 450 µl of Ric-8A and GTPγS in a quartz fluorescent cuvette prior to addition of 50 µl of 10 µM mGαi1 for a final concentration of 0.5 µM Ric-8A, 1 µM mGαi1, and 10 µM guanosine 5’-O-[gamma-thio]triphosphate (GTPγS) in 500 μl. GTPγS binding to size exclusion chromatography purified complexes of Ric-8A: ΔN31Gαi1 with and without nanobodies were compared to intrinsic nucleotide exchange of ΔN31Gαi1. GTPγS was added to preincubated complexes for final concentrations of 1 μM complex (or ΔN31Gαi1) and 10 μM GTPγS. For each assay, a minimum of 5 technical replicates were taken for each sample; more replicates were performed if the series of assays was conducted over several days, to control for changes in sample activity. Progress curves were fit to single or double (If a slow binding phase was detected) exponential rate models using SigmaPlot 7. Statistical significance of rate differences between reference and test samples was determined by a two-tailed Student’s t-test. Probability that differences are derived from a random distribution is reported. All data points are shown in box plots that show mean and standard deviation for each data set^12^.

### Crystallization of the ΔN31Gαi1:Ric-8A:3Nb complex

Crystallization trials for the Ric-8A:ΔN31Gαi1:Nb8109:Nb8117:Nb8119 complex were conducted by vapor diffusion using commercially-available crystallization screening kits. Sitting drops were set on 96-2 well INTELLI-PLATEs (Art Robbins Instruments) using a Gryphon crystallization robot (Art Robbins Instruments) at 3-10 mg/mL protein complex at a 1:1 v/v ratio with precipitation solution. Initial crystallization conditions were identified from hits on the ShotGun screen (MD1-88 Molecular Dimensions). Further optimization was carried out by grid screening variations in sodium malonate (Hampton Research) concentration and pH and by crystal seeding by hanging drop on 24 well VDXm plates (Hampton Research). Crystal seed stocks were prepared with the Seed Bead kit (Hampton Research) in 1.4 M sodium malonate pH 6.9. 0.9 µl of protein stock was added to 0.6 µl of reservoir and 0.3 µl of crystal seed stock and incubated at 12 °C for a minimum of 1-2 weeks. Optimal crystals were obtained from hanging drops containing 3.6 mg/ml Ric-8A:ΔN31Gαi1:3Nb, 1.4 M sodium malonate pH 6.9 at 12 °C.

### Crystallographic data collection and processing

Diffraction data were measured at the NSLS-II FXS beamline from LN_2_ cryoprotected crystals measuring approximately 50μ X 50μ X 5μ employing a helical data collection mode^13^, using X-rays of 0.9793 wavelength. A Si (111) double crystal monochromator was used to focus the X-ray beam to a spot size of 1μ X 1μ at the sample. Data were measured in the Phi axis rotation mode in 0.2° fames, at 0.06s/frame at a beam attenuation of 0.7. Diffraction data were measured on an Eiger 16M detector at a 133 Hz frame rate. Due to significant (> 0.5Å) differences in unit cell axis dimension between crystals, data from a single crystal were selected for data processing and structure determination. In view of the considerable anisotropy of diffraction (approximately 4.6Å along *a*\* and *b\** and 3.3Å along *c\**), three separate data processing strategies were conducted using the AutoPROC v1.0.2 software toolbox^14^ for evaluation in subsequent model-building and refinement steps. All three utilized XDS^15^ for data indexing and initial integration. The Standard Isotropic protocol uses SCALEA and TRUNCATE from the CCP4^16^ to generate isotropic data with a resolution cutoff (4.6Å) determined by the criteria R_pim_ ≥ 0.6, I/σ(I) ≥ 2.0 and CC_1/2_ ≥ 0.3. The Extended Isotropic protocol employed POINTLESS and AIMLESS scaling and analysis software^17^ to generate scaled intensities extending to the 3.2Å resolution with I/σ(I) ≥1.2. The Anisotropic Filtering protocol implemented in STARANISO^18^ generated a dataset that incorporates intensities within a locally averaged value of I/σ(I) to define an anisotropic diffraction cut-off surface (**Extended data Table 2)**.

### Crystallographic model building and refinement

All crystallographic calculations were conducted using the PHENIX 1.1.6 software package^19^ unless otherwise noted. Initial crystallographic phases for the Ric-8A:ΔN31Gαi1:3NB complex were determined by Molecular Replacement using atomic coordinates of Ric-8A (residues 1-426; PDB ID:6NMG), the Ras domain of Gai1:GDP (PDB ID: 1BOF), and an anti-VGLUT nanobody (PDB ID:5OCL) as search models. An initial atomic model, consisting of coordinates for Ric-8A residues 1-426, residues 34-65 and 190-320 of ΔN31Gαi1 and three nanobody backbone models were manually refit into a sigma-weighted 2mFo-DFc map using Coot v0.8.6^20^ and refined using phenix.refine. Completion and refinement of the crystallographic model followed the following general strategy: 1) model-fitting to the cryo-EM reconstruction in regions of the structure in which the path of the polypeptide chain was not defined in (or differed substantially from) the 2mFo-DFc map followed by real space refinement with secondary structure and geometry restraints using phenix.real_space_refine^21^; 2) refitting of the latter to the 2mFo-DFc map and subsequent refinement; 3) refinement of the cryo-EM map (see below) using the crystallographic model as an alignment reference. In the regions in which the polypeptide path observed in the cryo-EM-derived and crystallographic models diverged, no attempt was made to bring them into agreement. Initial cycles of crystallographic model building and refinement were carried out using Standard Isotropic or Extended isotropic data sets derived from datasets derived from merging and scaling data from the two most strongly diffracting crystals. The final rounds of fitting and refinement of the complete model (excluding the C-terminus of Gα1, the Helical Domain, and Switch I, residues 51-184) utilized an Anisotropic Extended dataset comprising data from the single crystal that afforded the strongest diffraction and optimal merging statistics (see **Extended Data Table 2**). A TLS model was applied during refinement using phenix.refine. Model quality and correlation with the refined electron density were performed using MolProbity^22^. The atomic coordinates for the refined crystallographic model for the Ric-8A:ΔN31Gαi1:3Nb complex and associated structure factors are deposited in the RCSB Protein Data Bank^23^ (PDB ID 6TYL). Figures depicting atomic models were rendered using PyMol version 2.3 (Schrodinger, LLC)

### Cryo-EM data collection

Three microliters of the Ric-8A:ΔN31Gαi1:4Nb complex at 0.4 mg/ml with 0.01% NP40 were applied onto glow-discharged 200-mesh R2/1 Quantifoil grids (Electron Microscopy Sciences). Grids were blotted for 4 s and rapidly cryocooled in liquid ethane using a Vitrobot Mark IV (Thermo Fisher Scientific) at room temperature and 100% humidity. The samples were screened using a Talos Arctica cryo-electron microscope (Thermo Fisher Scientific) operated at 200 kV and then imaged in a Titan Krios cryo-electron microscope (Thermo Fisher Scientific) with GIF energy filter (Gatan) at a magnification of 130,000× (corresponding to a calibrated sampling of 1.06 Å per pixel). Micrographs were recorded using EPU software (Thermo Fisher Scientific) with a Gatan K2 Summit direct electron detector, where each image is composed of 30 individual frames with an exposure time of 6 s and a dose rate of 11.5 electrons per second per Å^2^. A total of 8,670 movie stacks were collected with a defocus range of −1.5 to −3.5 µm.

### Cryo-EM image processing

All micrographs were motion-corrected using MotionCor2^24^ and the contrast transfer function (CTF) was determined using CTFFIND4^25^. All particles were autopicked using the NeuralNet option in EMAN2 v 2.31 and further checked manually, yielding 768,736 particles from selected 8,468 micrographs. Particle coordinates were then imported to Relion v3.0.6^4^, wherein multiple rounds of 2D classification were performed to remove poor 2D class averages. Meanwhile, ∼ 20,000 particle images were selected to build an initial model using the “ab-initio 3D” program in cryoSPARC v2^26^. A total of 676,130 particles were used for 3D classification in Relion to remove the no-conforming particles. Next, five rounds of 3D heterogeneous refinement, CTF refinement were performed in cryoSPARC to further remove the poor 3D classes. The final 3D refinement with 327,493 particles was performed in cryoSPARC using the “non-Uniform refinement” option with a soft mask of the complex density and a 3.85Å resolution map was obtained. The resolution of the final map was estimated according to the 0.143 FSC criterion. A 3.9Å low-pass filter was to the final 3D map for better display (See more information in **Extended Data Fig. 2 and Table I**).

### Cryo-EM model building

The cryo-EM model for the Ric-8A:ΔN31Gαi1:4Nbs complex was constructed with a refined crystallographic Ric-8A:ΔN31Gαi:3Nb model (in which only the GTPase domain was modeled) and a model of the helical domain of Gαi1 in complex with Nb9156 from the crystal structure of Gαi:Nb9156 complex (unpublished data). The final Ric-8A:ΔN31Gαi1:4Nb cryo-EM model was rebuilt after refinement of the cryo-EM reconstruction using a Ric-8A:ΔN31Gαi1:4NB mask as described above, with multiple rounds of manual refitting of the cryo-EM density map optimized refinement using phenix.real_space_refine with secondary structure and geometry restraints^21^. Model quality was evaluated with MolProbity^22^.

### Small Angle X-ray Scattering data collection, analysis and modeling

SAXS was performed at BioCAT beamline 18ID at the Advanced Photon Source, Chicago, with in-line size exclusion chromatography (SEC-SAXS) to separate sample from aggregates and other contaminants. Protein sample (approximately ∼5 mg/ml in 50mM HEPES, pH 8.0, 150mM NaCl and 1mM TCEP) was loaded onto a Superdex 200 Increase 10/300 GL column, which was run at 0.7 ml/min in an AKTA Pure FPLC (GE Healthcare Life Sciences). After detection by an in-line UV monitor, the sample passed through the SAXS flow cell, a 1.5mm ID quartz capillary with 10µm walls. Scattering intensity was recorded using a Pilatus3 1M (Dectris) detector which was placed 3.67m from the sample affording access to a range of momentum transfer (*q*) from 0.0065Å^-1^ to 0.35Å^-1^ [*q*=4πsin(θ)/λ]. Exposures of 0.5 s were acquired every 2 seconds during elution. Data were reduced using BioXTAS RAW v1.6.0^27^. Buffer blanks were created by averaging regions flanking the elution peak and subtracted from exposures selected from the elution peak to create the I(q) vs q curves used for subsequent analyses. *Ab initio* molecular envelopes were computed by the ATSAS v2.6.2 package^28^ programs DAMMIN^29^. Ten bead models were reconstructed in DAMMIF^30^, which were aligned and averaged in DAMAVER^31^ (Volkov, 2003). The Molecular envelope was visualized, and atomic models fit to molecular envelopes using Chimera v1.10.2^32^. The conformational flexibility of Ric-8A:ΔN31Gαi1 was modeled by coarse-grained fitting with respect to experimental SAXS data using SREFLEX program in the ATSAS software package^28^. Normal mode analysis was conducted with automatic determination of rigid body units. The final disposition of rigid body units after application of normal mode projections was determined by rigid body refinement with respect to the computed SAXS profile^33^. CRYSOL software from the ATSAS software package was used to model scattering profiles from atomic coordinates.

**Amino acid sequence alignments** were conducted using Clustal Omega^34^ *via* its web-server (https://www.ebi.ac.uk/Tools/msa/clustalo/).

### Data Availability

Coordinates of Ric-8A:ΔN31Gαi1:4Nb model from cryo-EM are deposited in the RCSB Protein Data Bank (PDB) with ID 6UKT. Coordinates of Ric-8A:ΔN31Gαi1:3Nb model from crystal structure are deposited in the PDB with ID 6YTL. The cryo-EM reconstruction is deposited in the Electron Microscopy Data Bank (https://www.ebi.ac.uk/pdbe/emdb/) with id EMD20812. The small angle X-ray scattering data for the Ric-8A:ΔN31Gαi1 complex is deposited in the Small Angle Scattering Biological Database (SASBDB)^35^ with accession code SASDG95.

## Acknowledgements

This research was supported by NIH R01GM105993 (S.R.S); NIH P41GM103832, R01GM079429, and S10OD021600 (W.C.); NSF 1738547 (T-C.M); NIH R01-GM088242 (G.G.T.); resources of Instruct-ERIC, part of the European Strategy Forum on Research infrastructures (ESFRI), the Research Foundation-Flanders (FWO) for their support to Nanobody discovery, and the Strategic Research Program (SRP) of the Vrije Universiteit Brussel (E.P. and J.S.). The Integrated Structural Biology Core at the University of Montana Center for Biomolecular Structure and Dynamics is supported by NIH COBRE award P20GM103546 (S.R.S.). We thank Drs. Wuxian Shi and Martin Fuchs at the National Synchrotron Light Source II (NSLS II) for assistance with data collection at the FMX (17-ID-2) beamline, supported by NIH grant P41GM111244 and the Department of Energy (DOE), KP1605010. Z.D. is supported by the SSRL Structural Molecular Biology Program by the DOE and by NIH grant P41GM103393. Small angle X-ray scattering experiments were conducted at the Advanced Photon Source, operated for the DOE Office of Science by Argonne National Laboratory under Contract No. DE-AC02-06CH11357 with support of NIH grant P41 GM103622 and 1S10OD018090-01 for purchase of the Pilatus 3 1M detector. We thank Dr. Baisen Zeng for assistance with GEF assays and SPR quantitation of Nb-Ric-8A binding.

## Author Contributions

L.J.M. prepared and crystallized the Ric-8A:Gαi1:3Nb complex, assisted with X-ray data collection, conducted mutagenesis studies and analyzed kinetic data.

K.Z. carried out image acquisition and processing, generated and refined cryo-EM reconstructions.

T-C.M. fit and refined atomic models to cryo-EM and crystallographic data and conducted SAXS analysis.

J.J. prepared and screened samples for cryo-EM data collection, and assisted with data analysis.

C. Y-H. Conducted crystallization experiments and assisted with protein preparations.

S.L. assisted with cryo-EM data collection and processing.

C.J.T. prepared samples for nanobody development and conducted initial nanobody characterization.

T.I.Z. advised on with X-ray data collection and processing.

S.T. and A.W. generated nanobodies.

G.G.T. provided general consultation on project.

J.S. oversaw nanobody development.

W.C. oversaw cryo-EM studies.

S.R.S. designed project, provided overall supervision, and wrote the manuscript with assistance from co-authors.

## Supplementary Information

Supplementary Information is available for this paper

**Extended Data Table 1.** Cryo-EM Data Collection and Refinement Statistics.

| Extended Data Table 1 |  |
| --- | --- |
| Cryo-EM Data Collection and Refinement Statistics |  |
| **Data collection and processing** | Ric-8A(491):Δ31NGαi1:4Nb |
| Microscope | Titan Krios |
| Voltage (kV) | 300 |
| Camera | Gatan K2 Summit |
| Pixel size (Å) | 1.06 |
| Total Dose (e-/Å<sup>2</sup>) | 69 |
| Defocus range (μm) | -1.5 to -3.5 |
| Number of micrographs | 8,670 |
| Number of initial particles | 768,736 |
| Symmetry | C1 |
| Number of final particles | 327,493 |
| Resolution (0.143 FSC, Å) | 3.90 |
| **Atomic model refinement** |  |
| Software | phenix.real_space_refine |
| Clashscore, all atoms | 8.79 |
| Poor rotamers (%) | 0.57 |
| Favored rotamers (%) | 99.43 |
| Ramachandran favored (%) | 84.24 |
| Ramachandran allowed (%) | 15.76 |
| MolProbity score | 2.15 |
| Bond RMSD (Å) | 0.006 |
| Angle RMSD (Å) | 1.005 |

**Extended Data Table 2.**
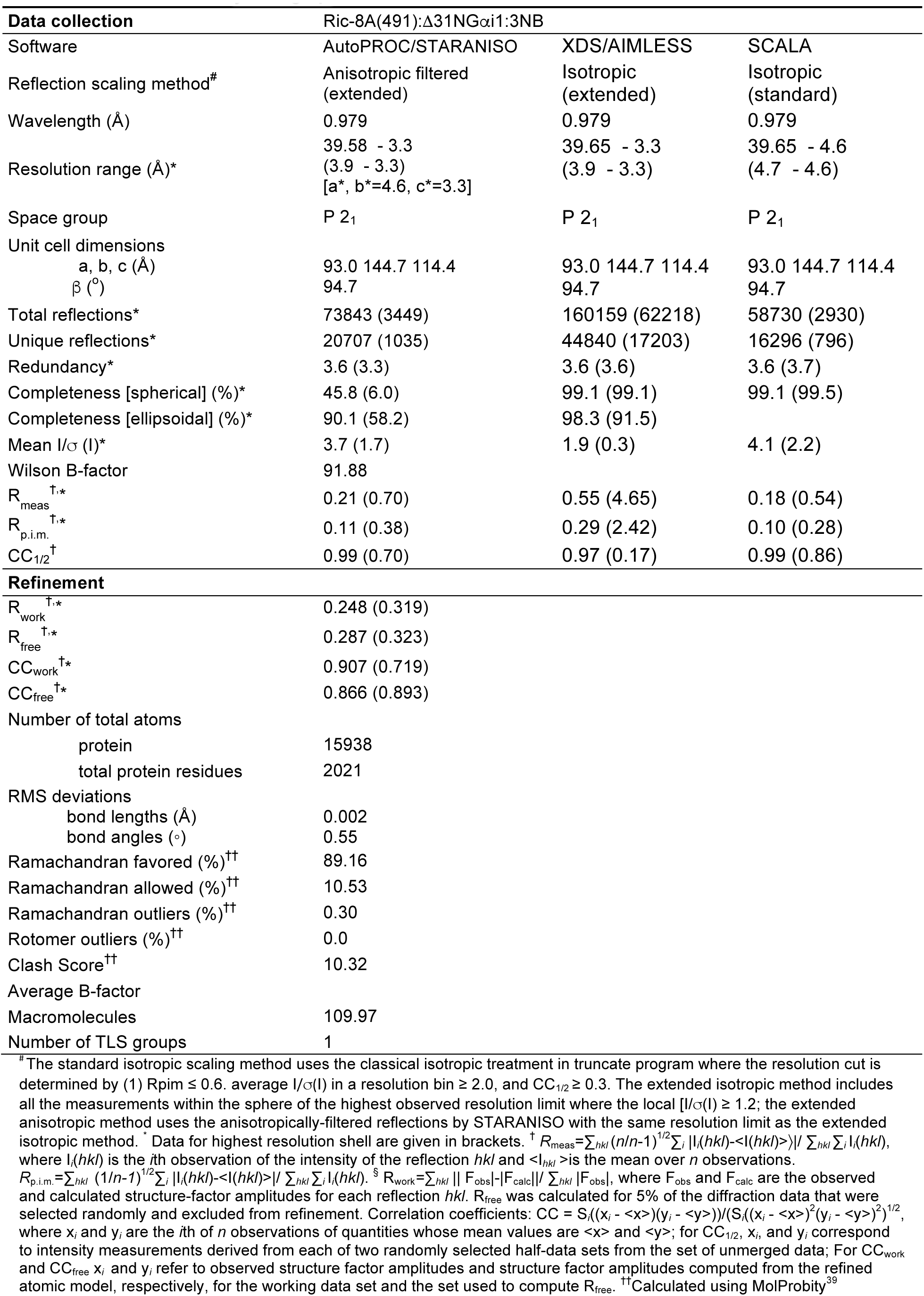
Crystallographic Data Collection and Refinement Statistics.

**Extended Data Figure 1.**
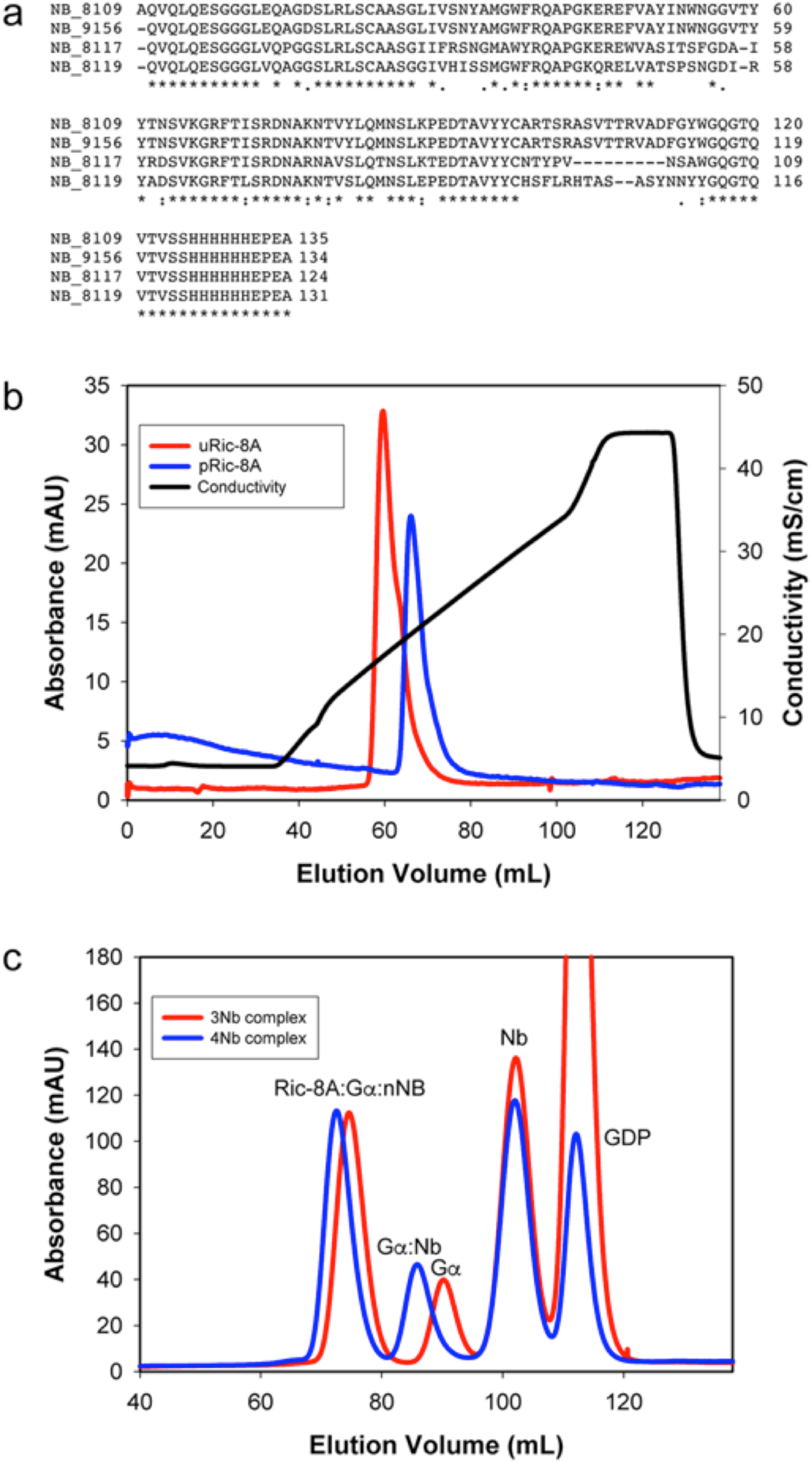
Purification of Ric-8A:Gα :Nb complexes. **a**, Amino acid sequences of nanobodies used to form complexes. **b**, Anion exchange chromatography of phosphorylated (pRic-8A) and unphosphorylated (uRic-8A). **c**, Gel filtration of complexes at the following molar stoichiometric ratios: 1 Ric-8A:2 Gα:2 Nb 8109:2 Nb 8117:2 Nb 8119 (*red)* and 1 Ric-8A:2 Gα:2 Nb 8109:2 Nb 8117:2 Nb 8119:4 Nb 9156 (*blue*) molar stoichiometric ratios, Ric-8A and Ga correspond to the 1-491 residue construct of Ric-8A and the 31N-terminal truncation mutant of Gαi1, respectively, as described in the main text. (see Methods).

**Extended Data Figure 2.**
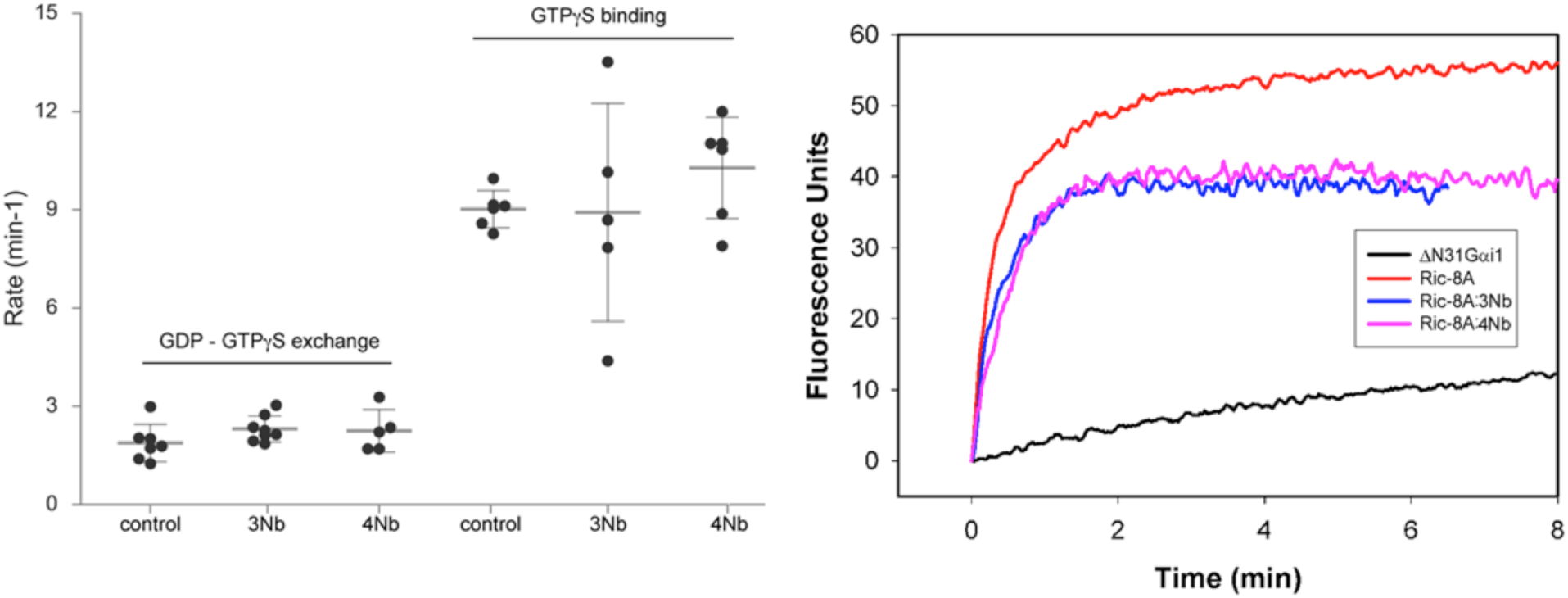
Effect of nanobodies on GEF activity of Ric-8A. **a**, GDP-GTP exchange rates were measured by the rate of tryptophan fluorescence increase upon addition of ΔN31Gαi1 to Ric-8A and GTPγS at final concentrations of 2mM Ric-8A, 1mM ΔN31Gαi1 and 10mM GTPγS in the absence (control) or presence of Nb 8109 (4 mM), Nb 8117 (4 mM), Nb 8119 (4 mM) (3Nb) or the latter with the addition of (4 mM) Nb 9156 (4Nb). GTP binding rates were measured by tryptophan fluorescence increase upon addition of GTPγS (10 mM final concentration) to size exclusion chromatography-purified Ric-8A:ΔN31Gαi1:nNb (1 mM). Progress curves for the 3Nb and 4Nb GDP-GTPγS exchange and 4Nb GTPγS binding assays were fit to a single exponential rate, whereas data for controls and the 3Nb GTPγS binding assay were fit to a double exponential model, due to the presence of a slow kinetic phase, and the kinetic constant of the fast rate (k1) reported. In all cases average and standard deviations of measurement are reported for a minimum of 5 experimental replicates. **b**, Sample progress curves for the data reported in panel **a**.

**Extended Data Figure 3.**
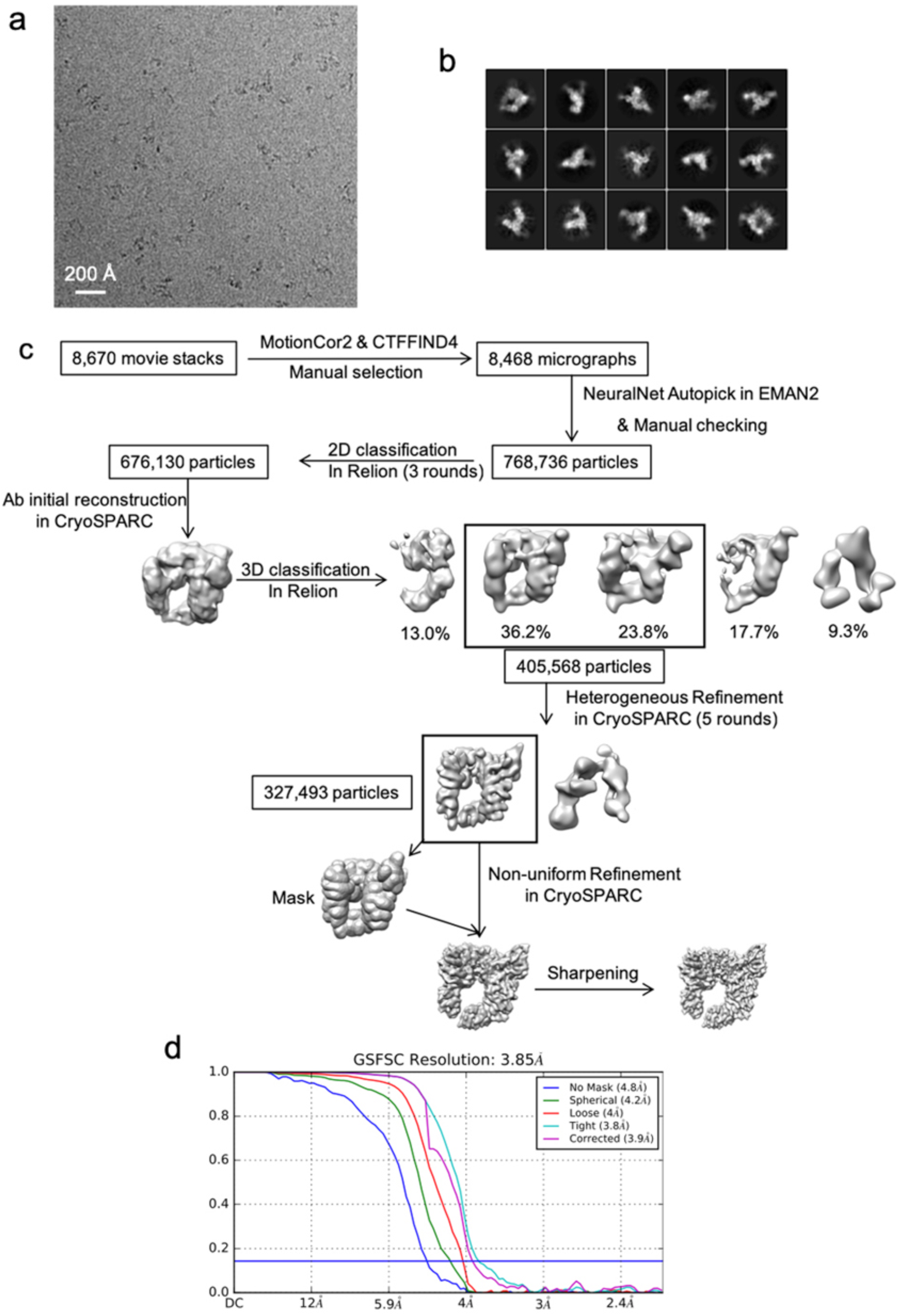
Single-particle cryo-EM of Ric-8A:Ga:4Nb complex and Work-flow of cryo-EM image processing. **a**, Representative motion corrected cryo-EM micrograph. **b**, Reference-free 2D Class averages computed in Relion. **c**, Work-flow for cryo-EM processing (see Methods). **d**, Gold-standard Fourier Shell Correlation plot for 3D reconstruction generated using cryoSPARK.

**Extended Figure 4.**
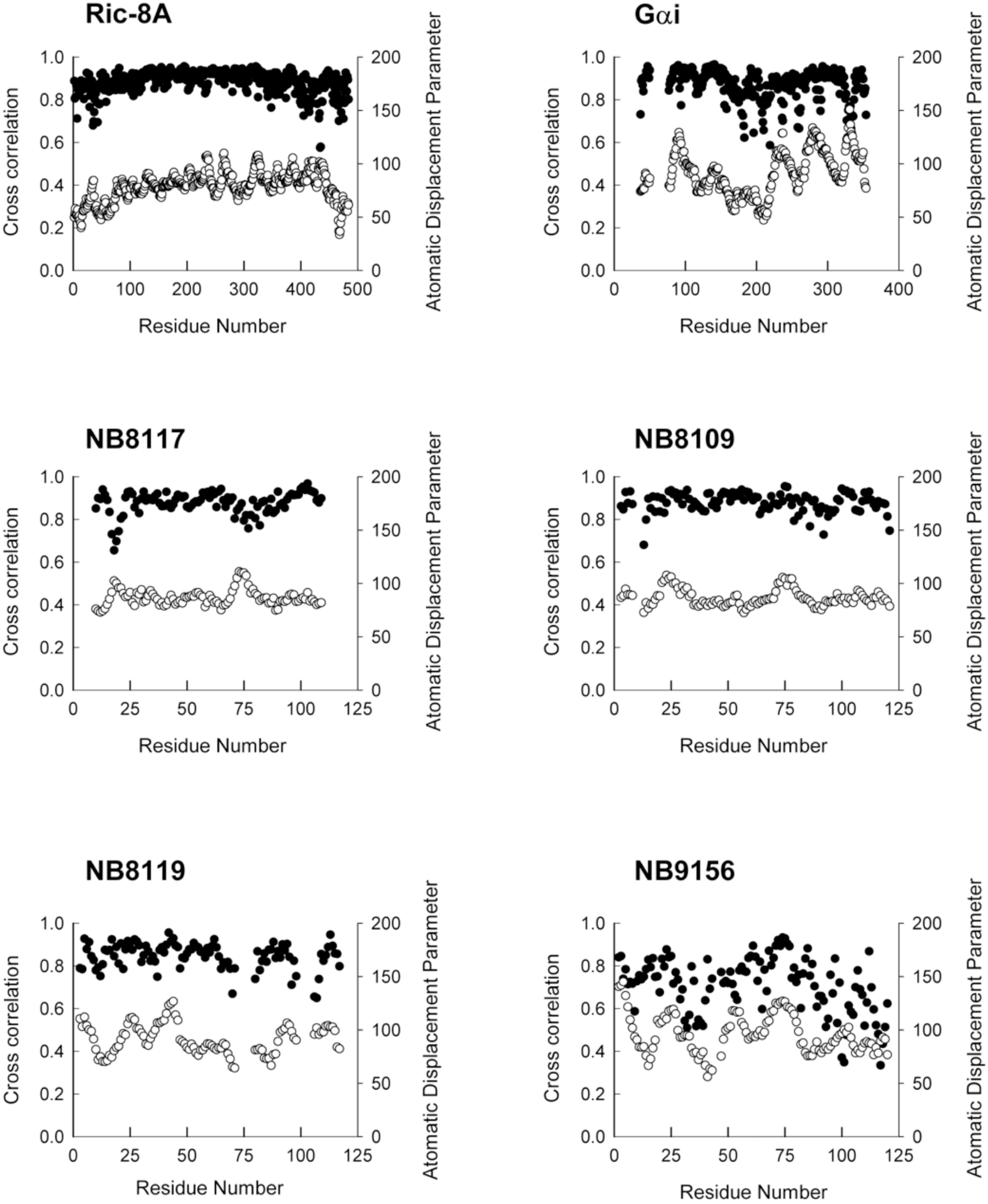
Real-space refinement of Ric-8A:Gα :cryoEM model. Model-map correlation (filled circles) and B factors (open circles) are plotted with respect to residue number for the macromolecular components of the Ric-8A:Gα:4NB model derived from cryo-EM. Data were generated using the phenix.real_space_refine program from the PHENIX suite.

**Extended Figure 5.**
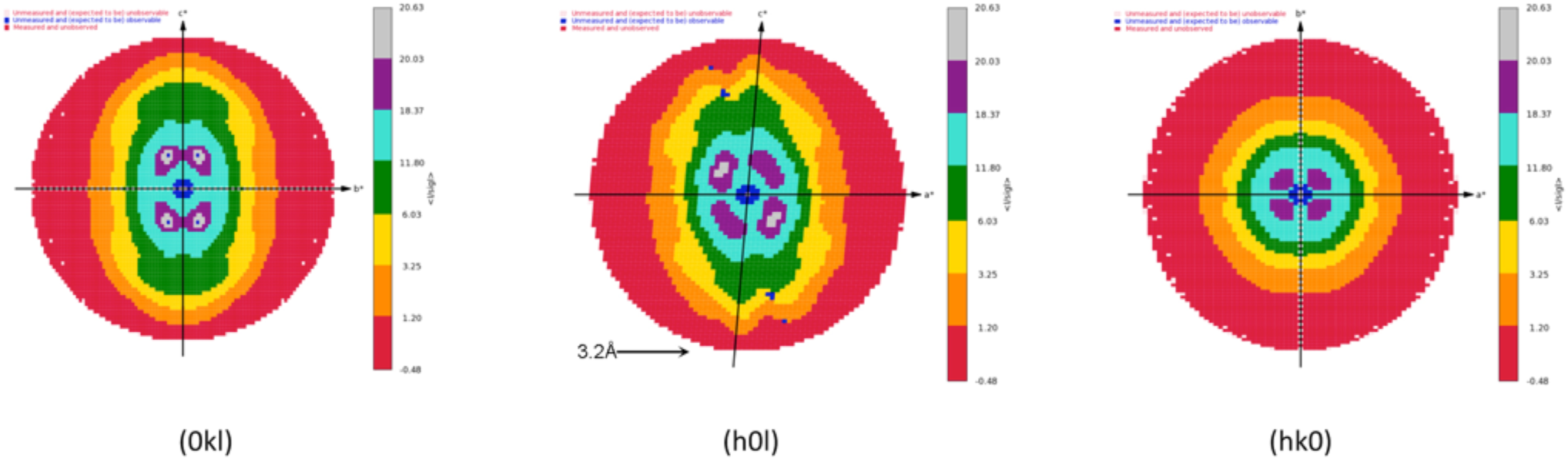
Projections of diffraction intensities from crystals of the Ric-8A:Gα :3Nb complex along reciprocal cell axes and color-coded by I/σI. Data generated using the STARANISO sever (see Methods).

**Extended Data Figure 6.**
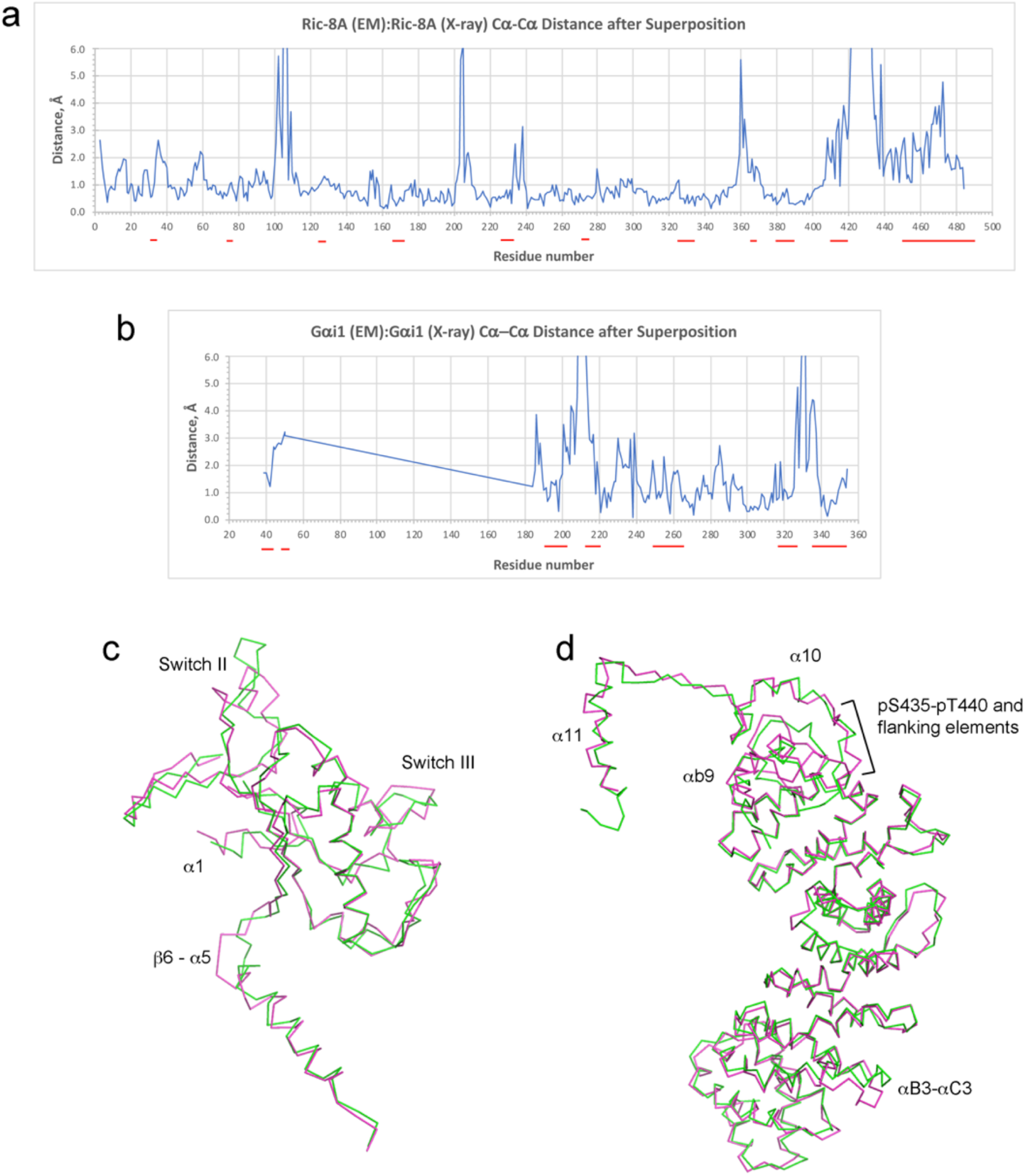
Agreement of cryo-EM and X-ray structures of Ric-8A:Gα i1. **a**, Deviation between corresponding Cα atoms after least-squares superposition of Ric-8A coordinates. *Red* lines below residue numbers mark regions at which Ric-8A contacts Gai1 (see also ED Figure 10). The RMSD for all 484 common Cα positions is 1.81Å. The PyMOL align routine yields an RMSD of 1.03Å after rejection of 69 outliers. **b**, Deviation between Gαi1 atoms, computed per panel **a**. The RMSD for all 183 common Cα positions is 2.36Å; PyMol align yields an RMSD of 1.49Å after rejection of 21 outliers**. c,** Superposition of cryoEM (*magenta*) and X-ray (*green*) structures of Gαi1. Structural elements that show the largest structural differences are labeled. **d**, Superposition of Ric-8A models, as per panel **c**.

**Extended Figure 7.**
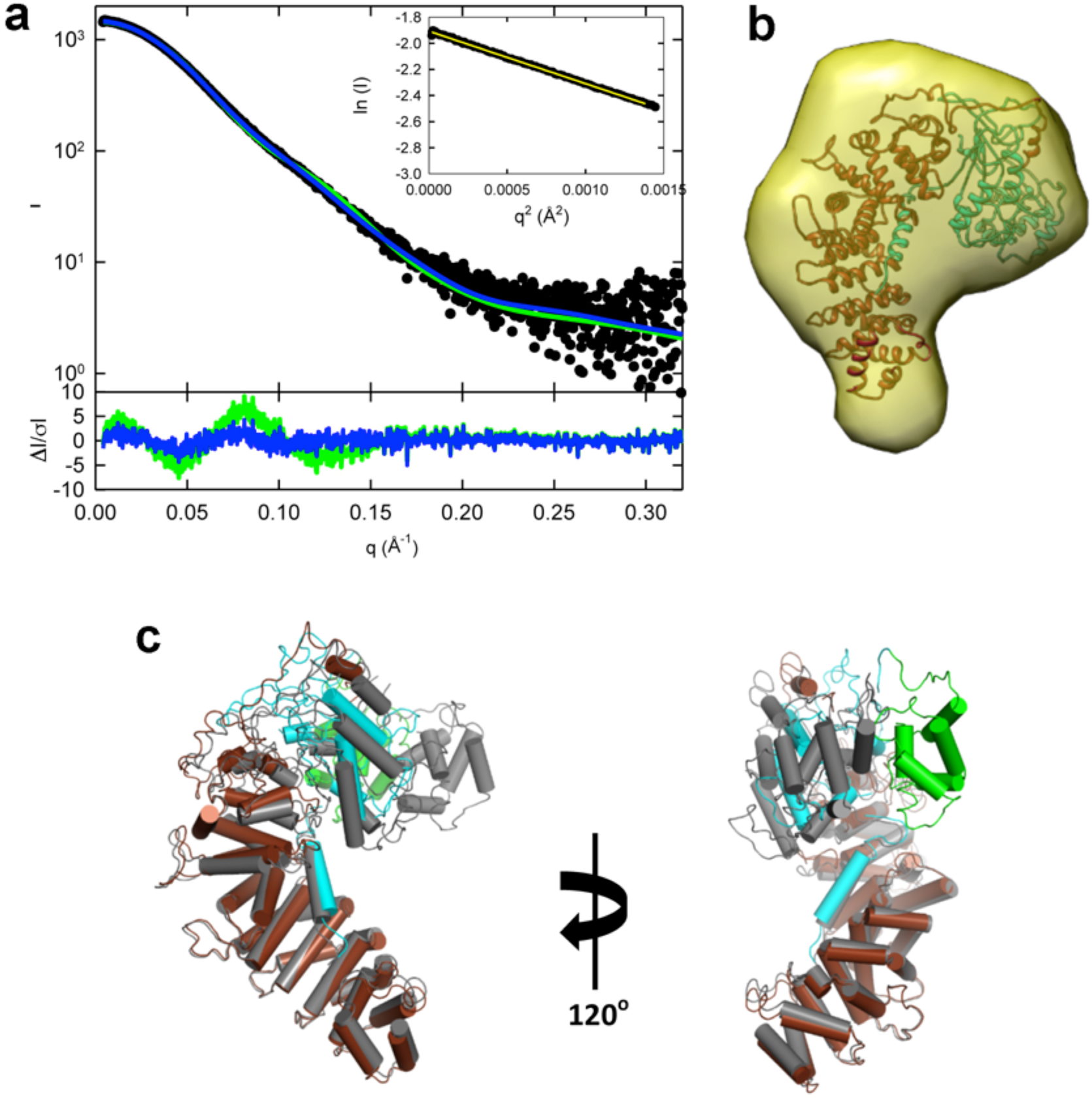
Solution structure and Normal Mode refined model of Ric-8A:Δ31NGαi1 complex from Small-Angle X-ray scattering. **a**, Solution scattering profile for Ric-8A:ΔN31Gαi1 complex (*closed circles*). Inset shows, associate Guinier plot. CRYSOL and SRELEX from the ATSAS software package were used, respectively, to compute scattering profiles from atomic coordinates, and use normal-mode analysis to generate a refined atomic model. Computed scattering curves for cryo-EM-derived atomic coordinates of Ric-8A:ΔN31Gαi1 complex are shown in *green* and that from normal mode analysis is rendered in *blue.* The χ^2^ for the fit of the computed scattering curve for the cryo-EM model is 5.48 and that for the normal-mode refined model is 1.24. The bottom panel shows the error-weighted residual difference plots ΔI/σI = (I(*q*)_experimental_ - CI(*q*)_model_)/σI(*q*) *vs q*, where C is a normalizing scale factor. **b,** SREFLEX-refined Ric-8A:ΔN31Gαi1 model fit to the *ab initio* molecular envelope calculated from the experimental SAXS scattering curve (*yellow*). **c**, Superposition of the SREFLEX-refined model of Ric-8A:ΔN31Gαi (Ric-8A, *brown*; GTPase domain of ΔN31GαI, *cyan*; Helical domain of ΔN31GαI, *green*) onto the cryo-EM model (*gray*), indicating a mode of helical domain rotation that may be preferred in solution in the absence of nanobodies. Models were superimposed using the Cα atomic positions of Ric-8A.

**Extended Data Figure 8.**
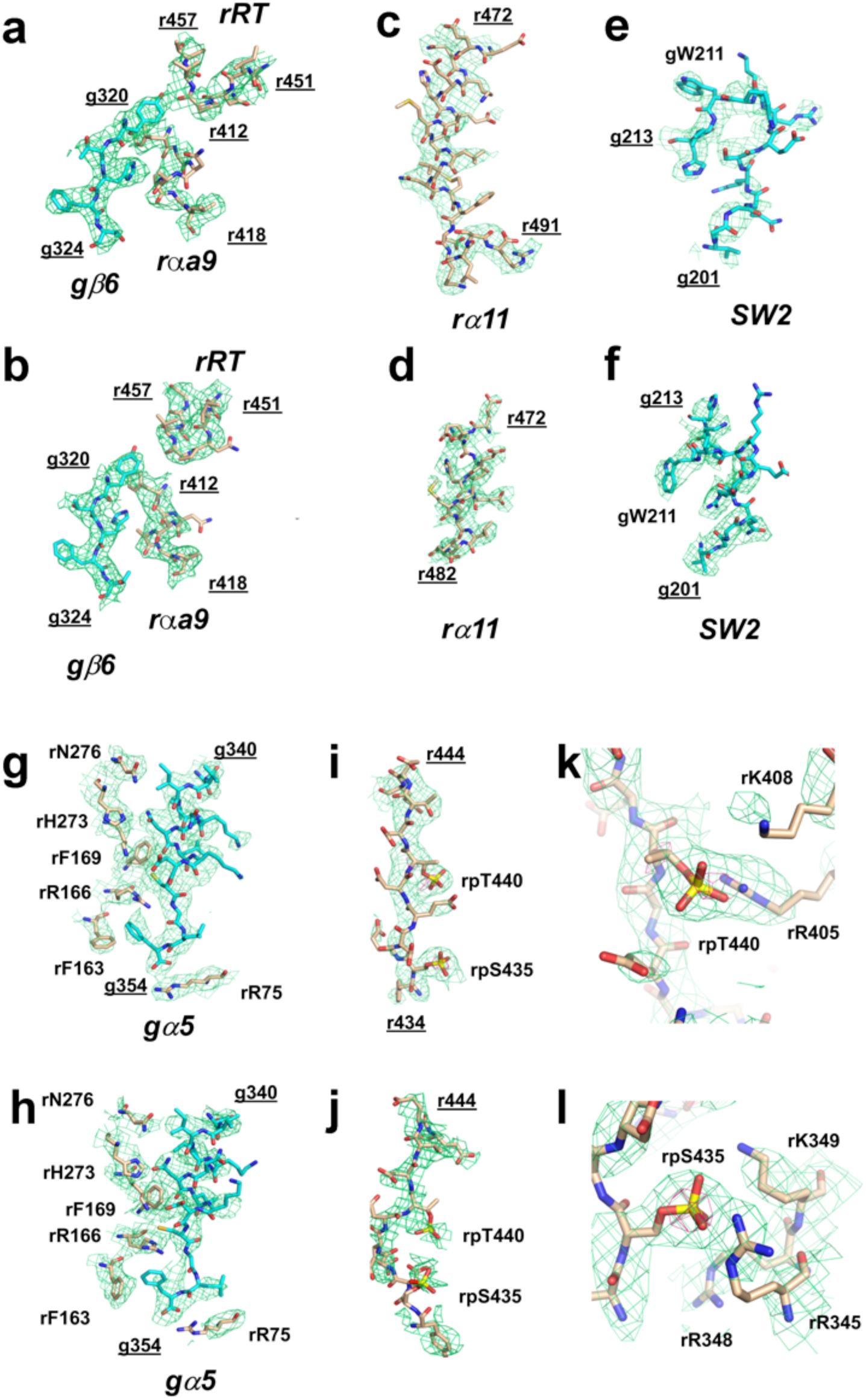
X-ray and cryo-EM density at selected residue ranges. 2DFo-mFc electron density is contured at 1.0 σ and cryo-electron density is contoured at a threshold of 3.5. Structural elements of Ric-8A and Gαi1 are labeled in bold italics, N- and C-terminal residue numbers of segments are shown underlined, selected residues are labeled. Carbon atoms in Ric-8A are colored *wheat* and in Gai1, *cyan*. Panels **a, c, e, g** and **i** show residues from crystallographic model with corresponding density. Panels **b, d, f, h** and **j** show the same from the cryo-EM model. Panels **k** and **l**, show the crystallographic model and associated density around phospho-serine 435 and phospho-threonine 440, respectively. The 5 s contour level is shown in *red*.

**Extended Data Figure 9a.**
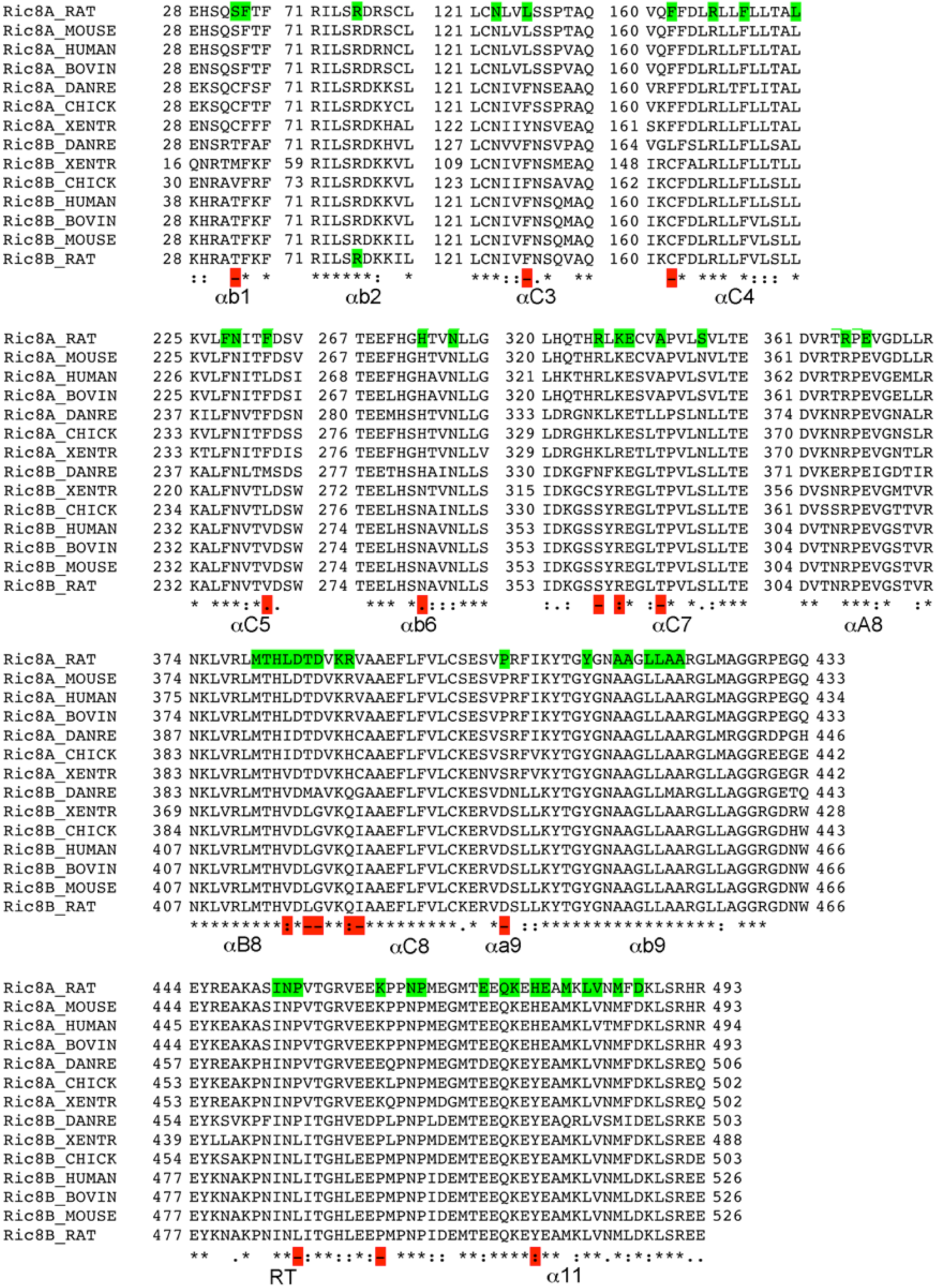
Amino acid sequences of Ric-8A and Ric-8B across phylogeny near Gα contact residues. Residue numbers are shown at the start and, for long tracts, the end of subsequences that harbor Ric-8A contacting-residues, highlighted in *green* in the sequence of rat Ric-8A, defined as those that include at least one atom within 4.0 of an atom in Gαi1. Amino acids not identical in rat Ric-8A and rat Ric-8B in *red* in the conservation summary at the bottom of each alignment. Central residues of secondary structural elements are shown below the conservation summary.

**Extended Data Figure 9b.**
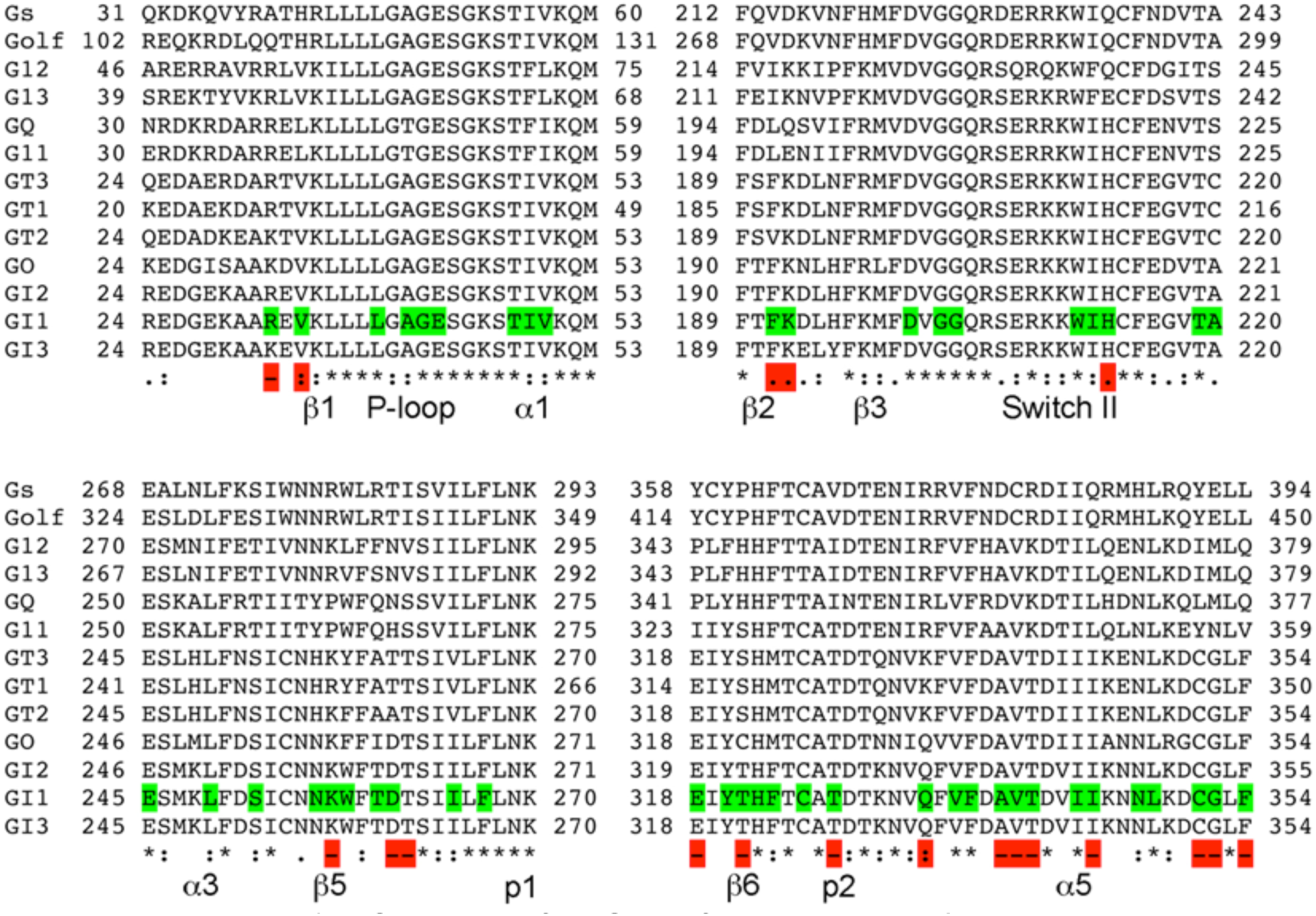
Sequence of rat Gα isoforms within Ric-8A contact regions. Residue numbers are shown at the start and end of subsequences that harbor Ric-8A contacting-residues, highlighted in *green* in the sequence of Gai1, defined as those that include at least one atom within 4.0 of an atom in Ric-8A. Amino acids not identical in Gαi and Gαs are highlighted in *red*, in the conservation summary at the bottom of each alignment. Central residues of secondary structural elements are shown below the conservation summary. Symbols p1 and p2 denote guanine nucleotide purine binding sites formed at the β5-αG and β6-α5 loops, respectively.

**Extended Data Figure 10.**
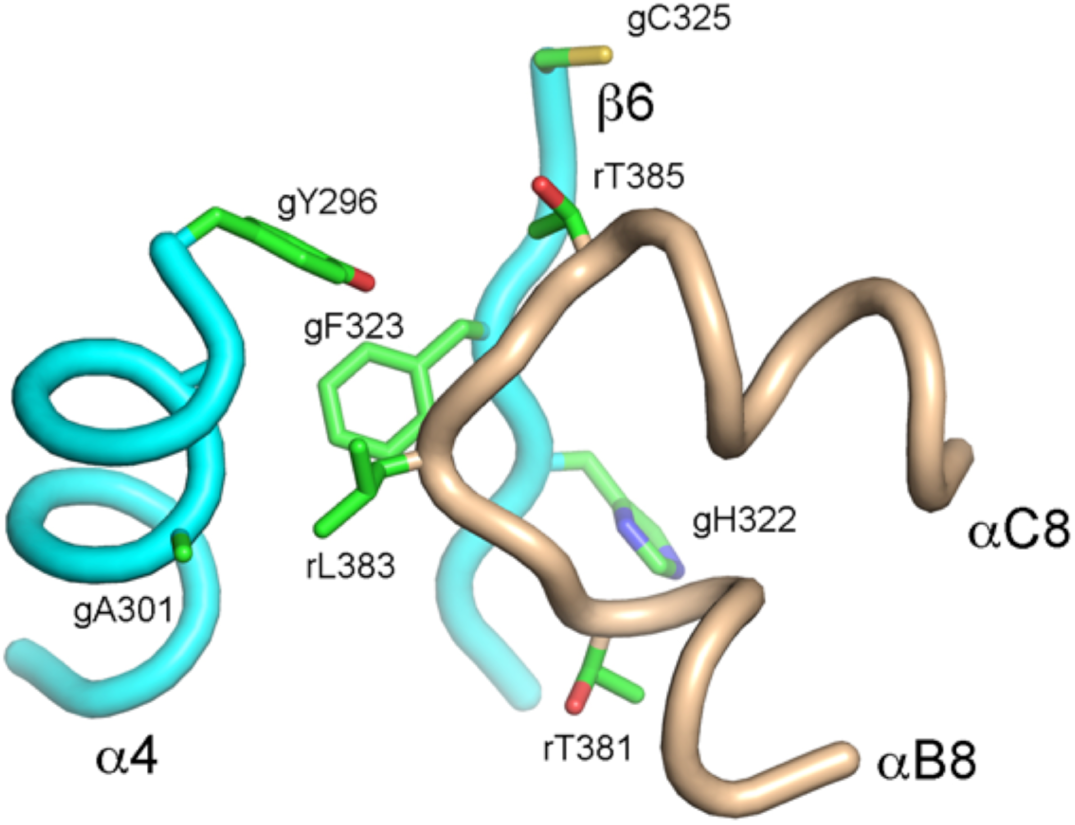
Detail of Ric-8A:Gαi1 contact involving Gα i1 α 4 and β 6 with Ric-8A αB8-αC8. The crystal structure is shown with Gαi1 backbone in *cyan* and Ric-8A backbone in *wheat* with side-chain atom colors, carbon, *green*, nitrogen, *blue*, oxygen, *red* and sulfur, *yellow*. Mutation of rL383 to glutamate reduces GEF activity to 50% that of wild-type Ric-8A.

